# Phloem-Limited Disease Impacts Citrus Carbohydrate Dynamics Distinctly from Source-Sink Manipulation

**DOI:** 10.64898/2026.10.19.760277

**Authors:** Dorcas Olufunke Alade, Jessica Gersony, Amit Levy, Christopher I. Vincent

## Abstract

Source-sink dynamics are defined by the interdependence of source and sink tissues and are key determinants of plant growth and productivity. This dynamic can, be altered by disease. One example is huanglongbing (HLB), a phloem-limited disease that impacts several economically important tree species by disrupting carbon partitioning and altering source-sink relationships. Though HLB is widespread in citrus, the specific mechanism in which phloem disruption impacts the co-regulation of carbon dynamics from source to sink has not been fully described. To address this, we took advantage of a natural study under field conditions, in which infected and uninfected trees were growing in the same field. After identifying trees that were uninfected, infected with symptoms, or infected without symptoms, we manipulated source:sink ratio by girdling branches and defoliating at 0%, 50%, and 75% of the original leaf area per fruit, while leaving a control treatment with no defoliation or girdling and used fruits as the sink of interest. We measured gas exchange, biomass, and non-structural carbohydrates in leaves, stems and fruits in response to these treatments. Our results show that leaf photosynthetic responses to HLB were primarily driven by stomatal regulation rather than biochemical limitations. HLB increased leaf dark respiration without altering the relationship between respiration and leaf soluble sugars. Among the plant organs, stem starch reserves were the most responsive to source-sink manipulation and were strongly correlated with multiple carbon pools and biomass in both fruits and leaves. Although HLB altered fruit carbon dynamics, it did not change fruit responses to source-sink manipulation. Overall, HLB affects source and sink tissues independently of the experimental source-sink manipulation. The stem serves as a transient carbon reserve and is sensitive to both HLB and source-sink manipulation. These findings raise further questions regarding total carbon fluxes and the importance of feedback inhibition of photosynthesis.

## Introduction

Plants grow, develop, and produce yield through coordinated source-sink relationships that regulate carbon production, transport, and utilization (Li *et al*., 2015; White *et al*., 2016). Source and sink activities adjust to developmental and environmental cues to meet the metabolic demands of growing tissues while maintaining efficient use of photosynthetically fixed carbon (Lemoine *et al*., 2013; Gessler and Zweifel, 2024). Source-sink coordination requires the integration of carbon assimilation in source tissues, phloem transport, and carbon utilization and storage in sink and transport tissues; and disruption at any point can alter whole-plant carbon balance and productivity (Khalilzadeh *et al*., 2026). Phloem-affecting diseases pose a major threat because they directly impair the vascular tissues responsible for source-sink transport (Bendix and Lewis, 2018).

In citrus, the phloem-restricted Gram-negative α-proteobacterium *Candidatus* Liberibacter asiaticus (*C*Las); the causal agent of huanglongbing (HLB; one of the most economically devastating disease of citrus) disrupts source-sink coordination (Singerman and Rogers, 2020; Cai *et al*., 2022; Zhao *et al*., 2025). HLB is transmitted by the Asian citrus psyllid (*Diaphorina citri* Kuwayama) (Tipu *et al*., 2021). *C*Las colonizes the phloem and induces structural impairments, such as callose deposition and sieve plates occlusion, that interferes with photoassimilate transport and disrupt source-sink balance (Achor *et al*., 2010; Gross *et al*., 2022). Despite evidence that HLB alters carbon transport and accumulation, little is known about how this disruption is expressed across the source-transport-sink pathway or how these tissues dynamically coordinate their responses.

Source-sink feedback allows plants to adjust photosynthesis according to carbon demand. Increased sink activity can stimulate photosynthesis by reducing carbohydrate accumulation, whereas sink activity reduce, starch builds up, which feeds back to inhibit photosynthesis by altering stomatal conductance and biochemical pathways (Syvertsen, 1994; Paul and Foyer, 2001; Ainsworth and Bush, 2011; Li *et al*., 2024). Thus, HLB would be expected to reduce photosynthesis and induce starch accumulation due to disrupted photoassimilate transport, while reducing sink growth, and consequently alter source-sink feedback responses. Yet the extent to which HLB modifies source-sink regulation of photosynthesis remains unclear.

Previous studies have reported reduced net photosynthesis and stomatal conductance, along with increased dark respiration, in HLB-affected citrus (Deng *et al*., 2019; Silva *et al*., 2021; Keeley *et al*., 2022; Khalilzadeh *et al*., 2026). Biochemical limitations may also contribute, particularly because HLB disrupts source-sink transport which could lead to feedback inhibition of photosynthesis. Similarly, the underlying cause of increased leaf dark respiration reported in HLB trees remains unclear. Elevated leaf respiration could represent a direct physiological response to HLB-associated defense and maintenance costs or an indirect response to increased carbohydrate availability. Distinguishing between these mechanisms is vital because each implies different physiological constraints and carbon costs, which may be relevant to other trees with similar diseases.

Transport tissues add another layer of complexity. Stems not only convey photoassimilates between the source and sink, but also a serve as dynamic carbon reservoirs that buffer fluctuations between carbon supply and demand (Furze *et al*., 2018). In perennial plants, stem nonstructural carbohydrates (NSCs) can modulate carbon balance when source or sink activity changes. Although, stems of HLB-affected citrus trees have been reported to accumulate more starch than those of healthy trees (Cai *et al*., 2022) suggesting that HLB may alter carbon storage or movement within transport tissues.

Although source-sink interactions are highly dynamic, with changes in one component eliciting compensatory responses in others. Existing research on HLB has largely considered source and sink responses independently (Baldwin *et al*., 2010; Tang *et al*., 2020; Cai *et al*., 2022; Keeley *et al*., 2022; Robledo *et al*., 2024). This limits our understanding of whether changes in leaf photosynthesis, stem carbon reserves, and fruit sink activity represent coordinated responses or independent consequences of disease. An integrated assessment of these tissues is needed to determine how HLB alters source-sink co-regulation and whether healthy and diseased trees differ in their responses.

Thus, we implemented a study to quantify source-sink co-regulation in healthy and HLB-affected citrus shoots using fruits as the sink of interest. We experimentally manipulated source-sink relationships using combinations of girdling and defoliation. We measured photosynthetic responses, leaf respiration, leaf, stem and fruit NSC dynamics, and examined relationships among carbon pools to determine how carbon supply and demand are coordinated across the source-transport-sink pathway. We hypothesized that: (1a) HLB-affected shoots would primarily limit photosynthesis through stomatal regulation, and HLB would mitigate photosynthetic upregulation in response to defoliation; (1b) elevated respiration in HLB-affected leaves is driven by increased substrate availability rather than by HLB-specific defense mechanisms; (2) stems of HLB-affected shoots would deplete carbon reserves in response to defoliation compared to healthy stems; and (3) fruits of HLB-affected trees would reduce sink activity more in response varying source:sink ratio relative to healthy trees. In addition, we aimed to determine whether NSC dynamics across leaves, stems, and fruits are coordinated to determine whether HLB-induced changes in carbon balance were expressed consistently across the source-transport-sink pathway. Our findings demonstrate important physiological characteristics of phloem-affecting disease and provide a framework that may inform the broader understanding of crop responses to similar pathogens.

## Materials and methods

### Study site and Plant material selection

The experiment was conducted using 8-years old *Citrus reticulata* ‘W. Murcott’ grafted on ‘US-897’ (*Poncirus trifoliata* [L.] Raf. × *C. reticulata* Blanco) rootstock. The orchard is a commercial grove located in Alturas, Central Florida, Florida, USA, characterized with Entisol soil that is about 80% Candler fine sand. Trees were grown in field soil under protective screens and were maintained under the standard irrigation regime used at the experimental site. Irrigation was provided through a micro-sprinkler system throughout the study.

With approximately 99% infection rate in mature trees in Florida, studies of HLB have been hampered by the inability to compare infected and uninfected trees under relevant environmental conditions (Graham *et al*., 2020). However, this study took advantage of a unique scenario. Trees were grown under protective screens that were temporarily breached during Hurricane Irma (2017), exposing a minority of trees to *Candidatus* Liberibacter asiaticus (*C*Las) while the rest remained shielded. The screen was repaired, and the psyllid vector population was eradicated, creating a field-like environment where a few trees were infected while the majority stayed healthy. HLB status was assessed through visual symptom evaluation to identify infected trees. This was then followed by quantitative PCR analysis of leaf, root, and fruit tissues to confirm the presence of *C*Las. Based on these evaluations, six citrus trees each of healthy and HLB-affected symptomatic and asymptomatic, were selected for the study (n=6).

### Identification and test for the presence of Candidatus Liberibacter asiaticus (CLas)

Diseased trees were first identified by visually examining the canopy and the visual identification was based on the following symptoms (Inoue *et al*., 2020; Tipu *et al*., 2021):

1. Leaves: asymmetrical blotchy pattern, vein corking.
2. Fruits: small, yellow-orange peduncular, green stylus, irregular shape.
3. Whole tree: shoot dieback, stunted or open canopy, off-season bloom.

Visual classification was then confirmed by quantitative PCR (qPCR) on leaf, root, and fruit tissues, and we observed consistent report from the fruit samples. Previous studies have consistently reported that DNA extracted from the fruit columella to capture the highest *C*Las loads (Kunta *et al*., 2013).

The qPCR *C*Las identification was done according to protocols reported in Levy *et al*. (2023)rom the columella, *C*Las DNA was extracted, pelleted, washed and resuspended in 100 μL of water for PCR analysis. qPCR was performed with primers and probes (HLBas, HLBr, and HLBp) for the ‘*C*Las’ bacterium using ABI PRISM 7500 sequence detection system (Applied Biosystems, Foster City, CA) in a 10-μL reaction volume consisting of the following reagents: 200 nM (each) target primer (HLBas and HLBr), 100 nM target probe (HLBp), and 5 × TaqMan qPCR Mix (Applied Biosystems). The qPCR assay consisted of 2 min incubation at 50 °C followed by 10 min incubation at 95 °C and 40 cycles at 95 °C for 15 s and 64 °C for 45 s, respectively. Data were analyzed and expressed as the DNA amplification cycle threshold (Ct) number.

Ct values were calculated based on a previously determined standard curve and we assigned qPCR Ct values ≤ 32 as positive for *C*Las infection (Levy *et al*., 2023). The data is presented in Table 2 with average Ct values of 35.36, 30.4 and 34.4 for healthy, HLB-asymptomatic and HLB-symptomatic respectively. qPCR-based CLas quantification showed that HLB-asymptomatic trees had the lowest Ct values, indicating the highest bacterial titer, while the Ct values of symptomatic trees were quite high. Previous studies have reported that trees with established HLB symptoms can still yield Ct values considered false negative due to the uneven distribution of *C*Las within the plant (Levy *et al*., 2023).

**Table 1:** List of abbreviations and variable names used in estimating gas exchange.

| Abbreviation | Variable | Units |
| --- | --- | --- |
| $V_{\text{cmax}}$ | Maximum carboxylation rate of rubisco | $\mu\text{mol CO}_2 \text{ m}^{-2} \text{ s}^{-1}$ |
| $J_{\text{max}}$ | Photosynthetic electron transport rate, associated with RuBP regeneration | $\mu\text{mol CO}_2 \text{ m}^{-2} \text{ s}^{-1}$ |
| $A_{\text{net}}$ | Net CO<sub>2</sub> assimilation | $\mu\text{mol m}^{-2} \text{ s}^{-1}$ |
| $R_{\text{dark}}$ | Dark respiration | $\mu\text{mol m}^{-2} \text{ s}^{-1}$ |
| $R_{\text{light}}$ | Daytime respiration | $\mu\text{mol m}^{-2} \text{ s}^{-1}$ |
| $g_s$ | Stomatal conductance to CO<sub>2</sub> | $\mu\text{mol CO}_2 \text{ m}^{-2} \text{ s}^{-1}$ |
| $L_s$ | Stomatal limitation; the reduction of $A_{\text{net}}$ by $g_s$ proportional to $A_{\text{net}}$ if $g_s$ were nonlimiting given $V_{\text{cmax}}$ and $J_{\text{max}}$ | |
| $g_s \text{ cost}$ | The reduction in $A_{\text{net}}$ imposed by $g_s$ | $\mu\text{mol CO}_2 \text{ m}^{-2} \text{ s}^{-1}$ |

**Table 2:**
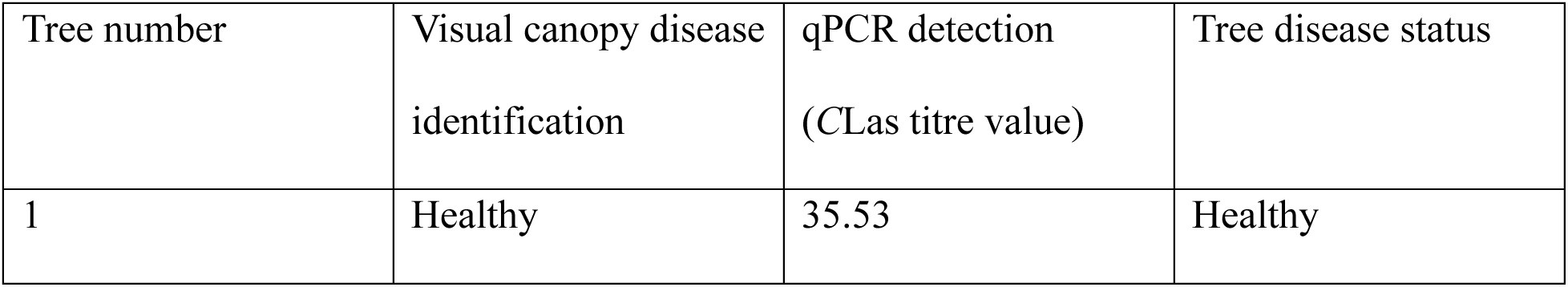
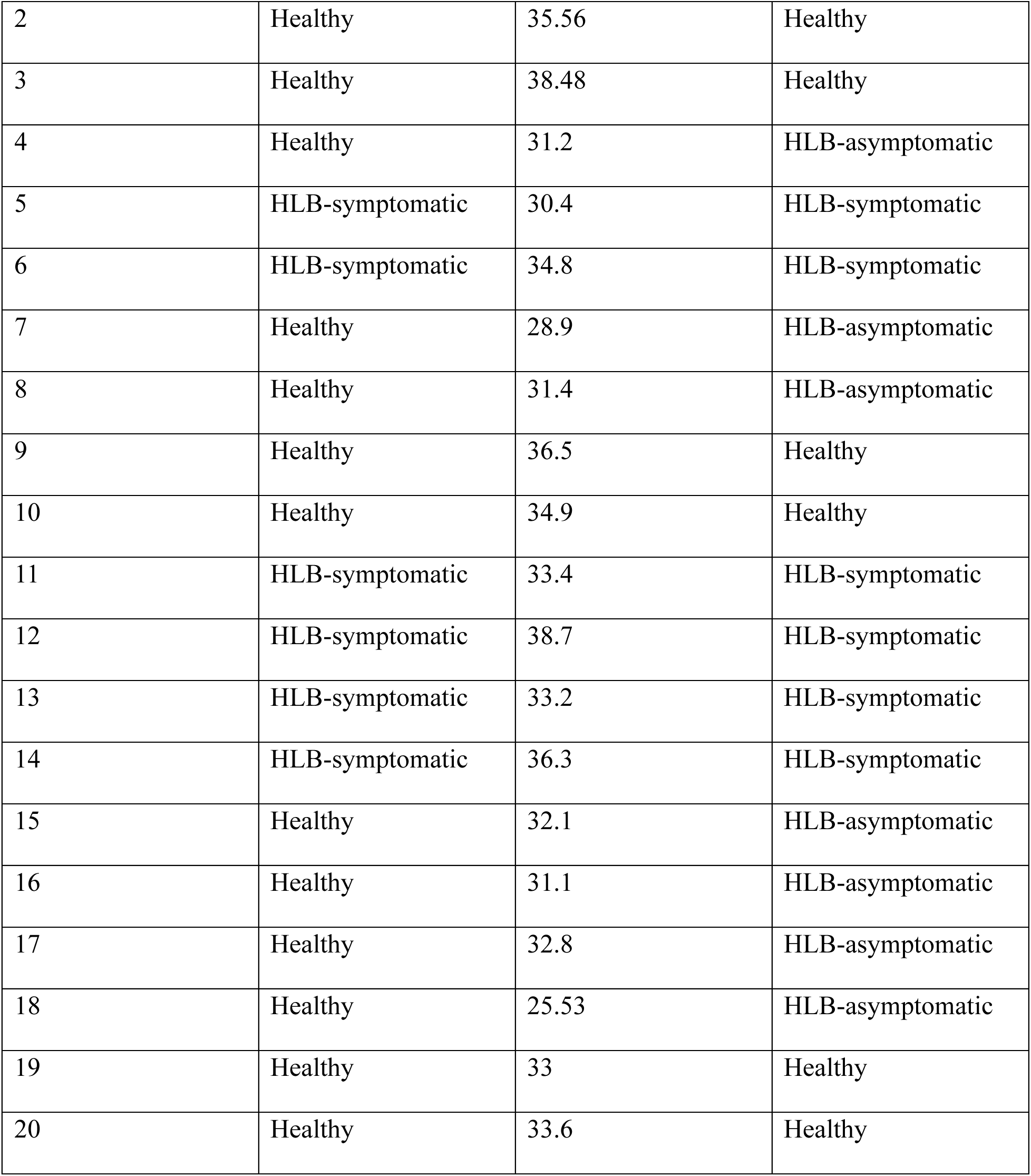
Tree *C*Las identification by visual and qPCR detection.

| Tree number | Visual canopy disease identification | qPCR detection (CLas titre value) | Tree disease status |
| --- | --- | --- | --- |
| 1 | Healthy | 35.53 | Healthy |
| 2 | Healthy | 35.56 | Healthy |
| 3 | Healthy | 38.48 | Healthy |
| 4 | Healthy | 31.2 | HLB-asymptomatic |
| 5 | HLB-symptomatic | 30.4 | HLB-symptomatic |
| 6 | HLB-symptomatic | 34.8 | HLB-symptomatic |
| 7 | Healthy | 28.9 | HLB-asymptomatic |
| 8 | Healthy | 31.4 | HLB-asymptomatic |
| 9 | Healthy | 36.5 | Healthy |
| 10 | Healthy | 34.9 | Healthy |
| 11 | HLB-symptomatic | 33.4 | HLB-symptomatic |
| 12 | HLB-symptomatic | 38.7 | HLB-symptomatic |
| 13 | HLB-symptomatic | 33.2 | HLB-symptomatic |
| 14 | HLB-symptomatic | 36.3 | HLB-symptomatic |
| 15 | Healthy | 32.1 | HLB-asymptomatic |
| 16 | Healthy | 31.1 | HLB-asymptomatic |
| 17 | Healthy | 32.8 | HLB-asymptomatic |
| 18 | Healthy | 25.53 | HLB-asymptomatic |
| 19 | Healthy | 33 | Healthy |
| 20 | Healthy | 33.6 | Healthy |

### Defoliation treatment

Four different branches bearing a single fruit were tagged on each tree to achieve four different defoliation ratios (Fig. 1). One of the branches was kept intact as control treatment (no girdling or defoliation), three were girdled to separate them from alternate sinks or sources and wrapped with parafilm to prevent the girdled part from desiccating. One of the girdled branches was left intact without defoliation; to represent high source:sink ratio and two of the girdled branches were defoliated to manipulate the source:sink ratio at 50%, and 75% of original leaf area, to represent low source:sink ratio. Treatments were applied at two developmental stages: early (treatment on May 18) and late (treatment on July 13) Stage II of fruit development which is characterized by rapid fruit enlargement driven by cell expansion (Matias *et al*., 2023). Data was collected between seven to eighteen days after defoliation treatment. At the end of each defoliation cycle, branches were destructively sampled for biomass and NSC assays.

**Figure 1:**
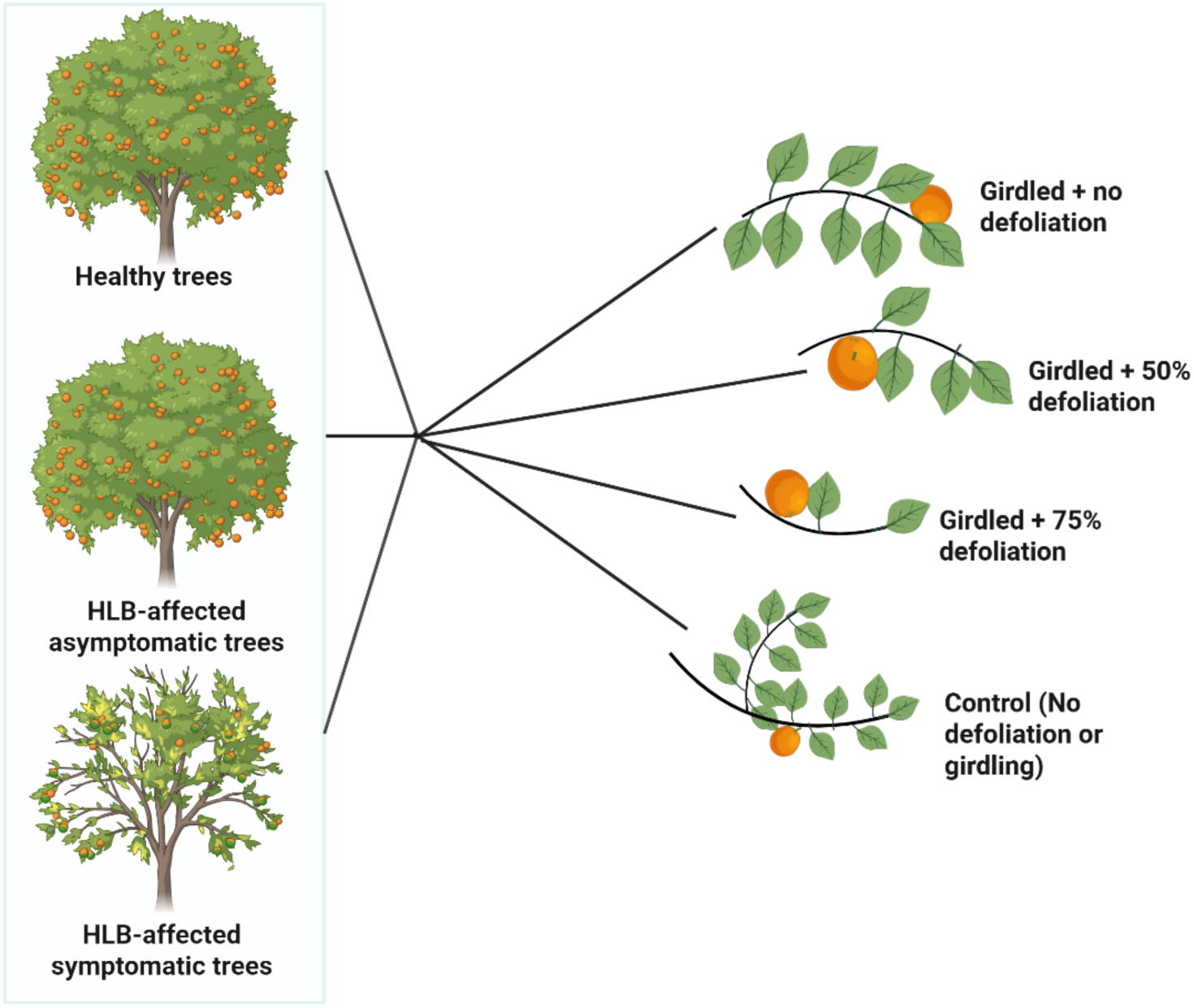
Schematic illustration of experimental conditions of source-sink manipulation (defoliation treatment) of healthy and HLB-affected ‘W. Murcott’ trees grown on field conditions under protective screens. For each HLB disease category four branches were selected for source-sink manipulation; Control (no girdling or defoliation), Girdled (no defoliation), Girdled + 50% defoliation and Girdled +75% defoliation.

### Gas exchange

#### Leaves

After one week of treatment, one fully expanded leaf per treated branch was measured within a single sunny day. To estimate carboxylation-related parameters, Rapid A/Ci response (RACiR) curves were measured between 10:00 and 12:00 hr on the leaf using an infrared gas analyzer (IRGA) LI-6800 (LI-COR Bioscience Inc., Lincoln, NE, USA). The environment settings used were (Li *et al*., 2024): flow rate of 300 μmol s^−1^, 0.1 kPa pressure, 30 mmol water, light intensity of 1200 μmol m^-2^ s^-1^, fan speed of 10000 rpm, CO_2_ concentration ramping from 100 to 900 μmol mol^−1^ at a rate of 100 μmol mol^−1^ min^−1^ and temperature was set according to the ambient temperature (28 – 30 °C). The minimum threshold of stomatal conductance to water used was 0.08 μmol m^−2^ s^−1^ ( Li *et al*., 2024). From the RACiR measurements, leaf photosynthetic activity: leaf *A*_net_ (leaf net CO_2_ assimilation), *V*_cmax_ (maximum rate of carboxylation), *J*_max_ (electron transport rate), *R*_light_ (day-time respiration), stomata limitation (*L*_s_) was estimated. On the same leaf used for RACiR measurements, dark respiration (*R*_dark_) was measured on a dark acclimated leaf (predawn) at environment settings used were flow rate of 300 μmol s^−1^, 0.1 kPa pressure, 30 mmol water, fan speed of 10000 rpm, light intensity of 0 μmol m^-2^ s^-1^, CO_2_ concentration of 400 μmol mol^−1^ and temperature was set according to the ambient temperature (28 – 30 °C).

#### Fruit

At the end of the experiment the fruit gas exchange was measured in a custom chamber attached to IRGA. The environment settings were as follows: flow rate of 600 μmol s^−1^, 0.1 kPa pressure, fan speed of 14000 rpm, 30 mmol water, CO_2_ concentration of 400 μmol mol^−1^, temperature was set according to the ambient temperature (28 – 30 °C). For fruit photosynthesis, measurements were taken on a sunny day between 9:00 AM and 2:00 PM under a light intensity of approximately 1000 μmol m^-2^ s^-1^ supplied to the chamber, while fruit respiration was measured under dark conditions.

### Nonstructural carbon (NSC) Sucrose and starch assay

Leaf, stem, and fruit samples were flash-frozen, stored at -20°C, ground and freeze-deied using a lyophilization. NSCs were measured using colorimetric methods as employed by (Kaur *et al*., 2023). Non-structural carbon was determined by extracting soluble sugars, sucrose and starch from 25 mg of dry leaf, stem or fruit ground samples. The samples were boiled after adding 500 μl of Milli Q (MQ) water for 10 mins at boiling point (100 °C) and agitated before centrifuging for five minutes at 13,000 rpm. After this the samples were separated into supernatants and pellets, the supernatant was used for reducing sugar and sucrose analysis while the pellet was used for starch quantification.

For the sucrose analysis, 50 μl of potassium hydroxide was added to 50 μl of the supernatant and the sucrose standard solution, the samples were boiled for 10 minutes to eliminate other reducing sugars except sucrose. The samples were cooled and then 1000 μl anthrone reagent was added to each tube. The samples were agitated and incubated for 10 minutes at 100 °C and shaken again after incubation. From each sample tube, 150 μl was pipette in duplicates into a 96-well plate and read using spectrophotometer at 620 nm wavelength.

For the soluble sugar analysis, 50 μl of the supernatant and the glucose standard solution were added into new tubes and 1000 μl anthrone reagent was added to each tube. The samples were agitated and incubated for 10 minutes at 100 °C and shaken after the incubation again. From each sample tube, 150 μl was pipette in duplicates into a 96-well plate and read using spectrophotometer at 620 nm wavelength.

The starch analysis was done by first washing off impurities by adding 1 mL of 100% ethanol to each pellet tube, then agitated and centrifuge for 10 minutes at 13,000 rpm. The supernatant was removed by decanting without disturbing the pellets, after which 1 mL MQ was added to the pellets, agitated and centrifuged for 10 mins at 13,000 rpm, and the supernatant was removed again gently without disturbing the pellets. Then 500 μl of 0.2M Sodium acetate buffer (pH 5.5), 100ul of 700 units/ml amyloglucosidase and 100ul of 70 units/ml alpha amylase were added for enzyme degradation, after which the samples were agitated and incubated at 37 °C for two hours. The samples were shaken after incubation, centrifuged at 13,000 rpm for five min. Thereafter, 50 μl of supernatant was handled according to the same anthrone-based protocol used for sucrose and soluble sugars.

Calibration curves on every plate gave linear equations for soluble sugar, sucrose and starch content in the plant organs.

### Biomass measurement

The fruit diameter was measured with a caliper before and after the defoliation treatment. After sampling, the branches were dried by lyophilizing and the leaf, stem and fruit mass were measured using a weighing scale.

### Data analysis

Data and statistical analysis were done using R software (R Core Team, version 4.3.1, 2023). RACiR data were processed using plantecophys package (Stinziano *et al*., 2019) and using the same package A/Ci curves were created. From the data, photosynthetic parameters; *V*_cmax_ (maximum rate of carboxylation), *J*_max_ (electron transport rate), *A*_net_ (leaf net CO_2_ assimilation), *R*_light_ (day-time respiration), were estimated using Fitaci() command in plantecophys package. Stomatal cost (*g*_s_ cost) was estimated by subtracting the *A*_net_ estimate for each sample when *C*_a_ was 420 μmol mol^−1^ and that when *C*_i_ was 420 μmol mol^−1^ (the value when *C*_a_=*C*_i_ is the value expected when leaf conductance is non-limiting). Stomatal limitation (*L*_s_) was considered as the stomatal cost relative to the *A*_net_ at nonlimiting leaf conductance.

The normality of the data was tested using Shapiro–Wilk Test, thereafter, the effect of disease status, source-sink manipulation and fruit development stage were analyzed on data using a generalized linear mixed-effects model with lmer() command from lme4 package (Bates *et al*., 2015). The study was completely randomized for HLB status (whole plot), while source-sink manipulation treatments were nested within trees (split-plot). Thus, tree number was used as a random effect to account for the appropriate nesting structure. Analysis of variance was performed using anova() command. For significant effects (*P*-value < 0.05), means separation was performed using emmeans() command in the multcomp package (Hothorn *et al*., 2008), also using a significance threshold of *P*<0.05. To distinguish the effects of girdling from defoliation, treatment effects were evaluated for variables showing a significant source–sink manipulation treatment effect or treatment × disease interaction or treatment × phase interaction, using Tukey-adjusted pairwise comparisons. The ungirdled control was compared with the girdled treatment with 0% defoliation to evaluate the effect of girdling independent of defoliation. Defoliation effects were evaluated by comparing girdled branches subjected to 0%, 50%, and 75% defoliation. Linear mixed-effects models were fitted with treatment as a fixed effect and tree identity as a random intercept. For variables showing significant treatment effects, estimated marginal means were obtained using the emmeans package in R.

To test for relationships among continuous variables, since some of the variables might covary ggpairs() command from GGally package (Schloerke *et al*., 2025) was used to view the overall interactions, from which linear regression models were generated for variables with significant correlations with or without disease factor using lm() command and the significant relationships and interactions were tested using the anova() command.

## RESULTS

### Leaf net photosynthesis response to HLB is driven by stomatal regulation

We found no significant effects on the Calvin cycle fluxes (*V_c_*_max_, *J*_max_; Supplemental Fig. S1, S2 and Table S1) at both fruit development stages assessed. However, *A*_net_ was impacted by the interaction between source-sink manipulation and HLB disease (*P* = 0.04, Fig. 2A), with no effect of fruit development stages. In healthy branches, girdling or defoliation did not alter *A*_net_. In HLB-asymptomatic branches, girdling reduced *A*_net_ but defoliation had no effect. In contrast, defoliation significantly affected *A*_net_ in HLB-symptomatic trees. Specifically, 50% defoliation increased *A*_net_ relative to non-defoliated girdled branches, whereas 75% defoliation reduced *A*_net_ to levels below those observed in non-defoliated girdled branches.

**Figure 2.**
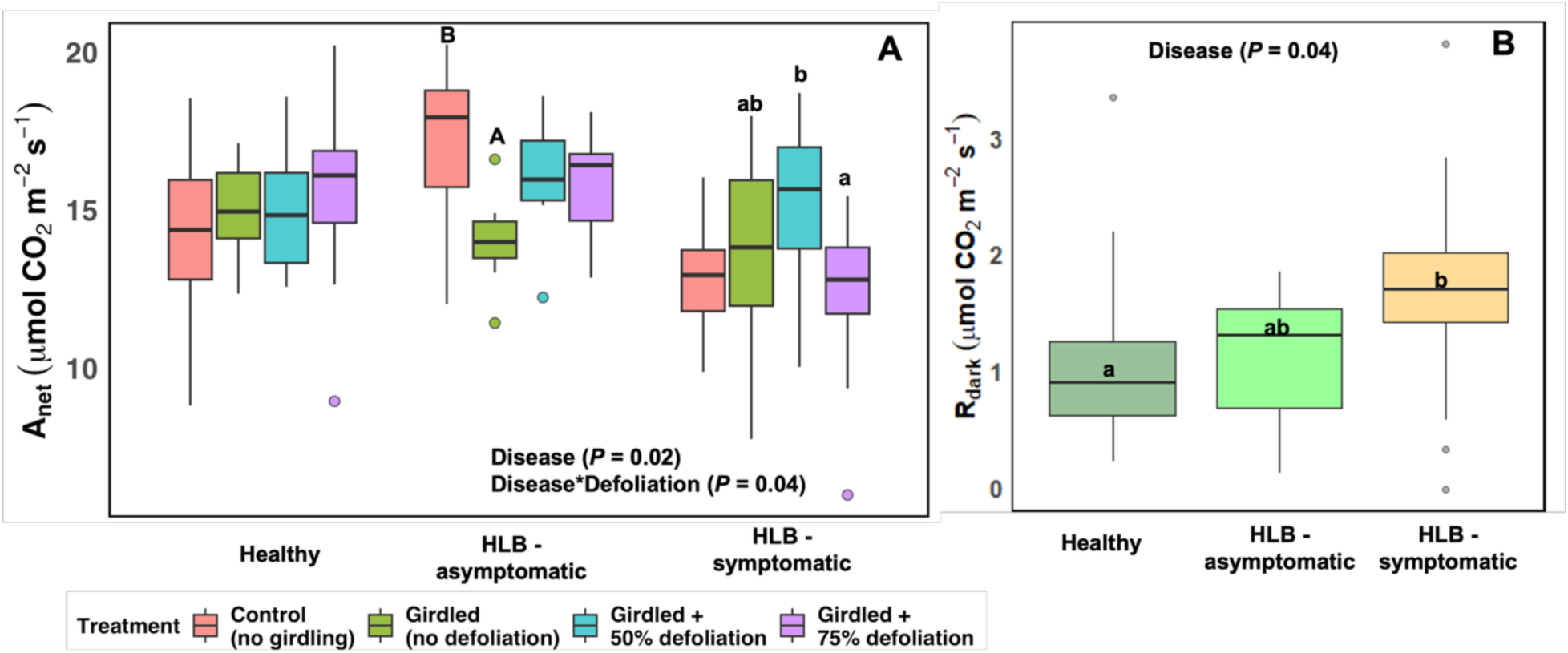
Leaf net photosynthesis (A; *A*_net_) and dark respiration (R_dark_; B) of *Citrus reticulata* ‘W. Murcott’ trees grown under protective screens in a study of Huanglongbing (HLB) disease and various source-sink manipulations. Data collected at sampling dates during Stage II of fruit development. Groups sharing different letters differ significantly (Tukey’s honestly significant difference test, *P* < 0.05). To distinguish girdling from defoliation effects, different capital letters indicate significant differences between the ungirdled control and girdled, non-defoliated treatment, whereas different lowercase letters indicate significant differences among defoliation treatments within each disease category.

*R_dark_* was not affected by source-sink manipulation or fruit development stages, but HLB increased *R_dark_* (*P* = 0.04, Fig. 2B). HLB-symptomatic leaves had greater *R_dark_* followed by asymptomatic leaves and healthy leaves.

Stomatal limitation (*L*_s_) represents the proportional effect of stomatal conductance (*g*_s_) on *A*_net_, given a specific *V_c_*_max_ and *J*_max_. *L*_s_ was significantly affected by the interaction of source-sink manipulation and HLB disease (*P* = <0.001, Supplemental Fig. S3), with no effect of fruit development stages. Leaves of HLB-affected symptomatic trees were the most responsive to *L*_s_, *L*_s_ was highest in control branches and was reduced when shoots were girdled, however, it increased drastically when 75% defoliated, suggesting a strong role for *g*_s_ in both HLB and in defoliation responses.

We observed distinct HLB disease effects on *L*_s_ in the control and 75% defoliation treatments (Fig. 3). Within control treatment (Fig. 3A), *L*_s_ showed only a possible effect (*P* = 0.07), however, when shoots were defoliated at 75% rate (Fig. 3B), HLB increased *L*_s_. Fig. 3C and 3D show the leaf CO_2_ exchange, including *A*_net_, *R*_dark_, and stomatal cost (*g*_s_ cost; the difference between *A*_net_ not limited by *g*_s_ and the measured *A*_net_ values). HLB disease did not change the stomatal cost in control treatments across disease categories, however at 75% defoliation, HLB significantly affect stomatal cost (*P* = 0.05; Supplemental Table S2). HLB affected *A*_net_ and *R*_dark_ in both control and 75% defoliated treatments. In the control treatment, *A*_net_ of healthy leaves were intermediate between HLB symptomatic and asymptomatic leaves. In 75% defoliation (3D), healthy and HLB-asymptomatic trees had similar *A*_net_, which was greater than HLB-symptomatic. In both cases, HLB-symptomatic had the lowest *A*_net_ and the highest *R*_dark_ and *g*_s_ cost.

**Figure 3.**
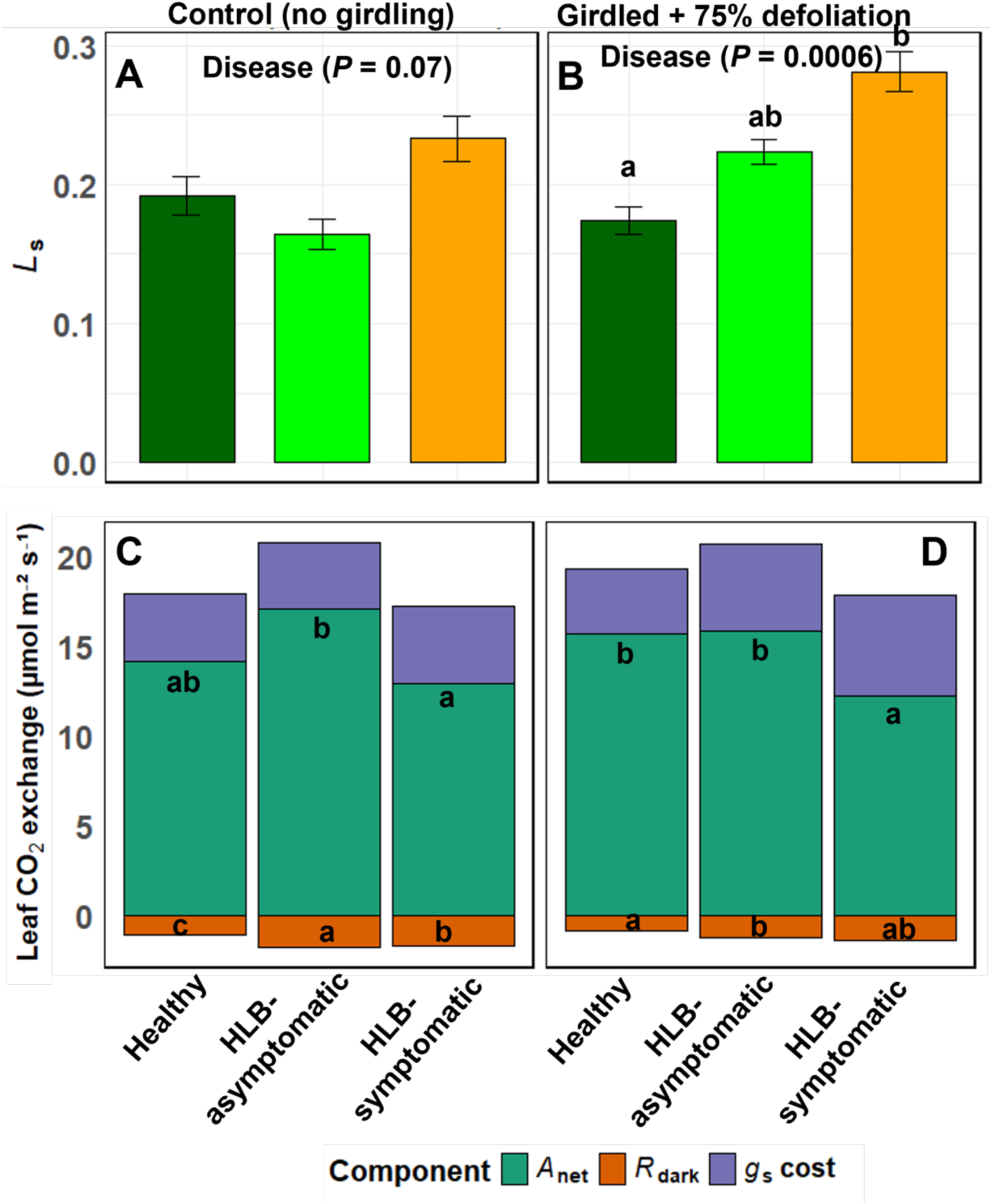
The effect of HLB-disease on stomatal limitation (L_s_; A and B) and leaf CO_2_ exchange (C and D) of field grown *C. reticulata* ‘W. Murcott’ trees under protective screens during Stage II of fruit development for Control treatment (A, C) and girdled treatment + 75% defoliation rate (B, D). Bars show means +/-standard error (n=6). Groups labeled with different letters are significantly different (Tukey’s Honestly Significant Difference; *P* < 0.05).

### HLB increased leaf R_dark_ and NSCs, which were correlated

To assess the effect of HLB on leaf NSC allocation and their relationship with leaf dark respiration, we measured leaf NSCs, including total NSCs, soluble sugar, sucrose, and starch (Table 3) and tested the correlations among these variables.

**Table 3:**
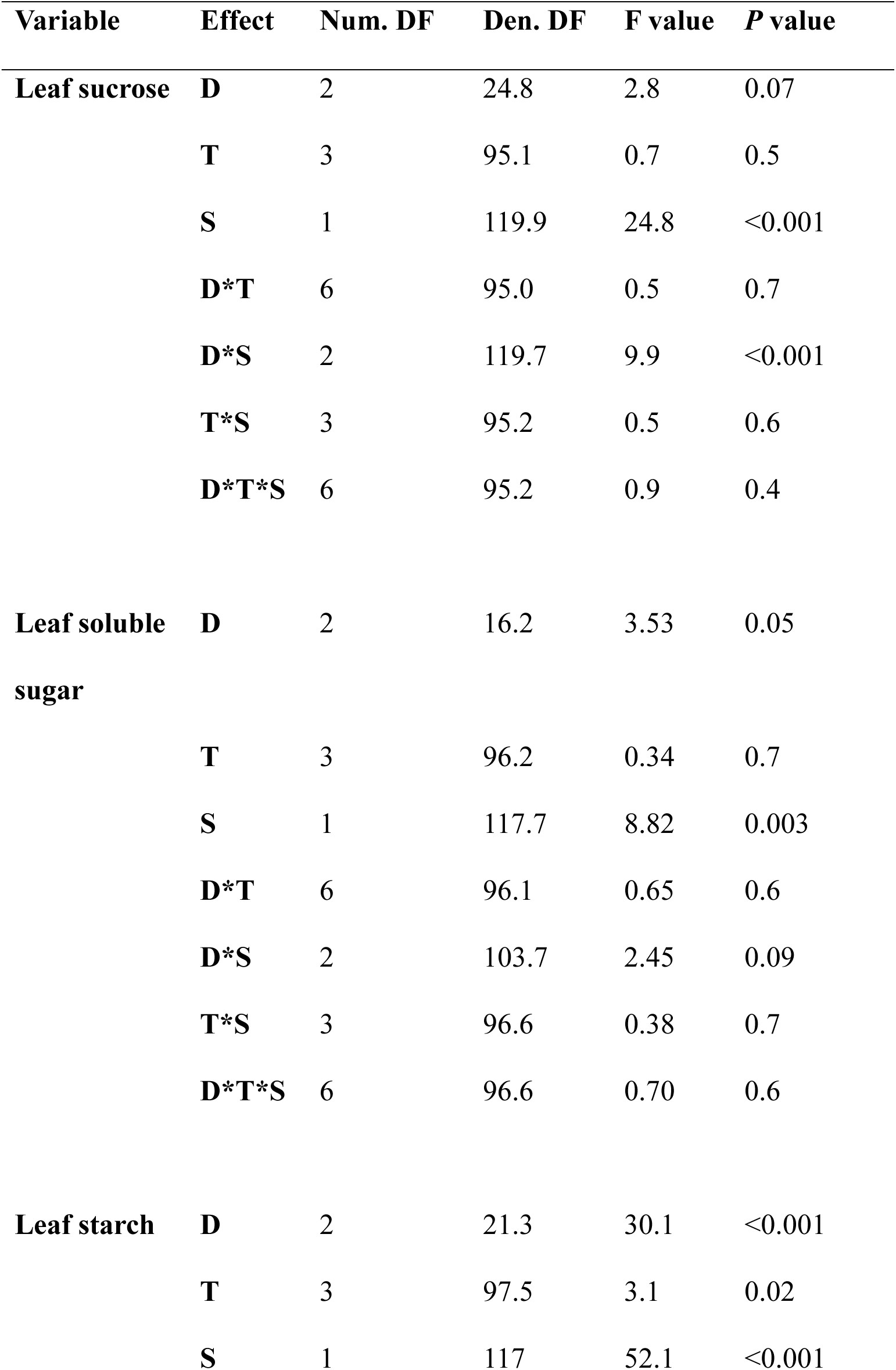
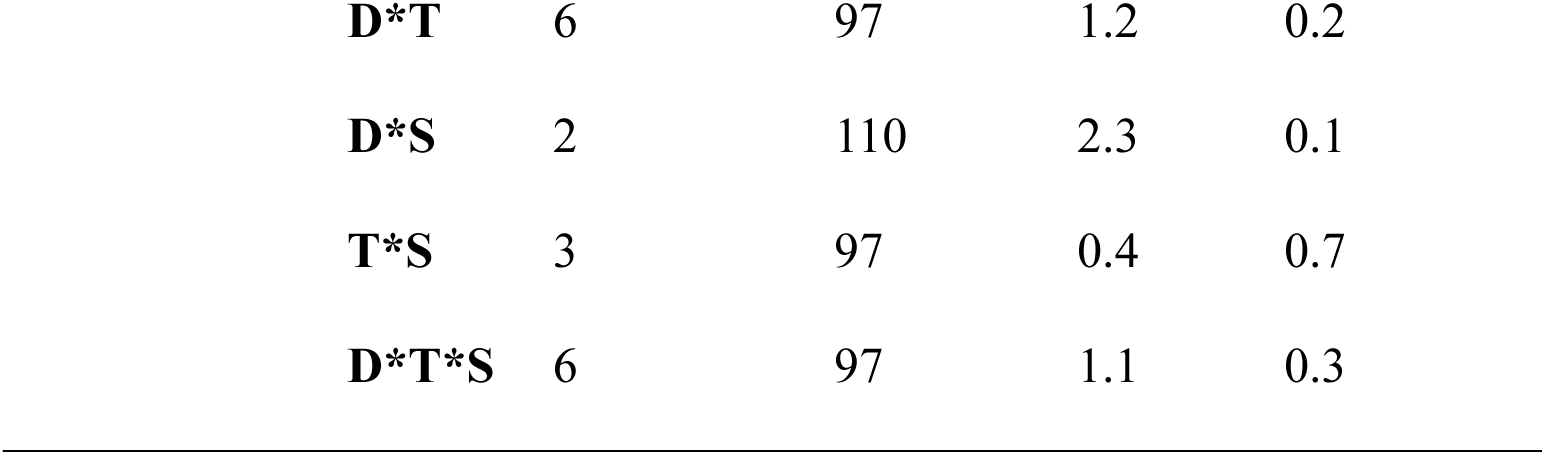
Analysis of variance of leaf non-structural carbohydrates of healthy and HLB infected *C. reticulata* ‘W. Murcott’ fruiting branches subjected to various source:sink manipulations. Source:sink manipulations included, untreated control, girdled, girdled and 50% defoliated, and gridled and 75% defoliated. D-Disease status; T-Source-sink treatment; S-Stages of fruit development.

While leaf sucrose content was not significantly impacted by source-sink manipulation, HLB disease showed a marginal effect on leaf sucrose content (*P* = 0.07, Supplemental Fig. S4). We observed an interaction between HLB disease and fruit development stage on leaf sucrose content (*P* < 0.0001, Supplemental Fig. S4). Specifically, at late Stage II, HLB-affected leaves had greater sucrose content than healthy leaves, even though overall leaf sucrose levels were greater in early Stage II than in late Stage II. Similar to the observed leaf sucrose response, HLB also had an effect on leaf soluble sugar content (*P* = 0.05, Supplemental Fig. S5).

HLB affected leaf starch content (*P* < 0.0001; Fig. 4A), with leaves of HLB-affected symptomatic trees accumulating more starch than healthy and HLB-affected asymptomatic trees. Source-sink manipulation also altered leaf starch content (*P* = 0.02): girdling increased leaf starch content, which then decreased upon defoliation (Fig. 4B). At late Stage II of fruit development, starch concentrations were greater than early Stage II (*P* < 0.0001; Supplemental Fig. S6) Correlations showed positive linear relationships between leaf carbohydrates content and leaf *R_dark_* (Fig. 5 A-C). *R_dark_* increased as total non-structural carbon (TNSC) (*P* = 0.001; r = 0.37), starch (*P* = 0.01; r = 0.3), and soluble sugars (*P* = 0.006; r = 0.3) increased. Based on these correlations, we tested linear regressions of *R_dark_* by leaf NSC, starch content and soluble sugar, including testing the effect of HLB and source-sink manipulation. Only HLB disease affected the intercept of the relationship though with low explanatory power, with *R*^2^ = 0.24; 0.20; and 0.26 for leaf NSC, starch content and soluble sugar respectively. There was no interaction effect between HLB disease and leaf carbohydrate contents on *R_dark_*, resulting in no effect of HLB on the slope of *R_dark_* response to any of the leaf carbohydrate contents. Interestingly, sucrose content did not significantly affect *R_dark_* (*P* = 0.3).

**Figure 4.**
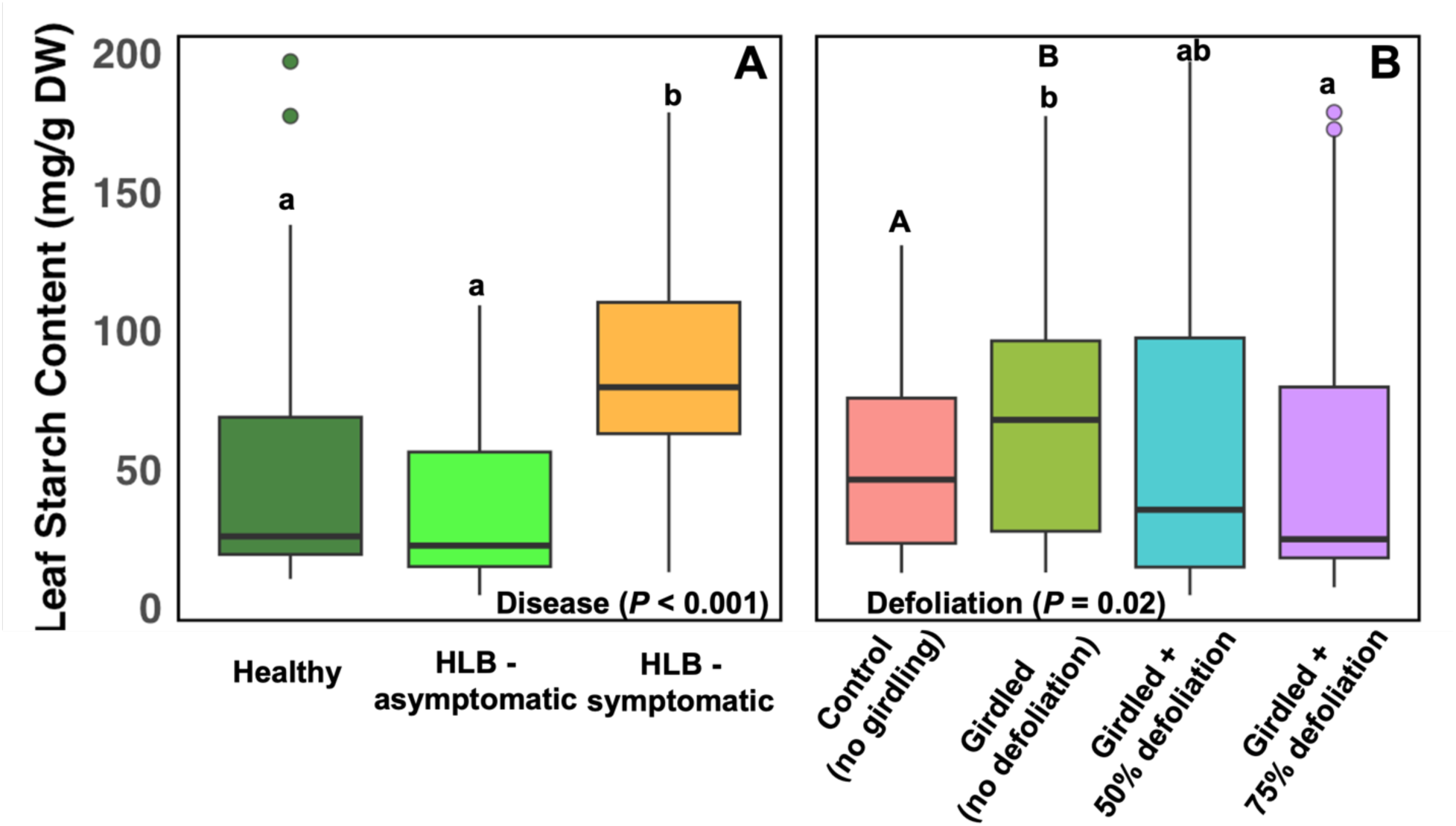
Leaf starch content response to HLB-disease status (A) and source-sink manipulation of all HLB disease categories (B) in *C. reticulata* ‘W. Murcott’ trees grown under field conditions under protective screens during Stage II of fruit development. Groups sharing different letters differ significantly (Tukey’s honestly significant difference test, *P* < 0.05). To distinguish girdling from defoliation effects, different capital letters indicate significant differences between the ungirdled control and girdled, non-defoliated treatment, whereas different lowercase letters indicate significant differences among defoliation treatments within each disease category.

**Figure 5.**
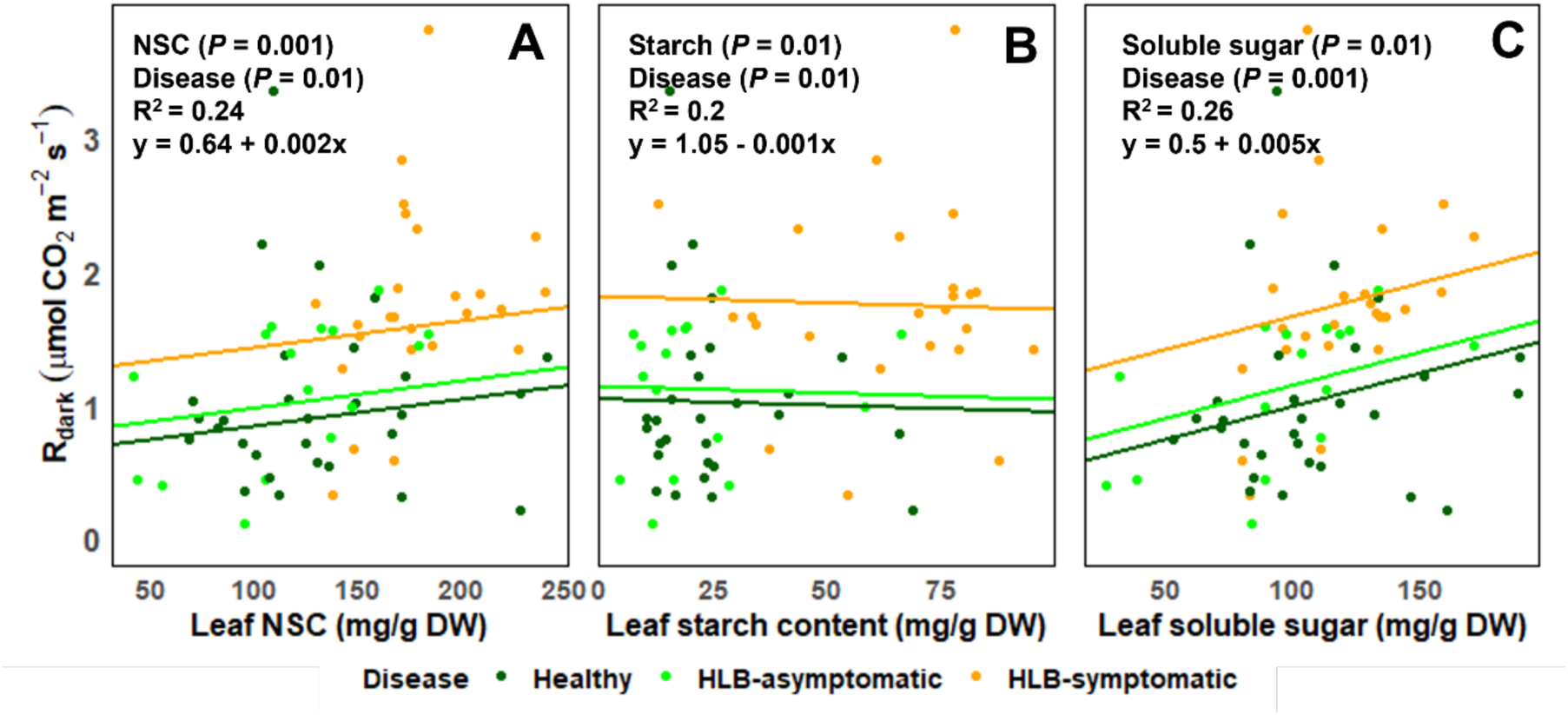
Leaf dark respiration (*R_dark_*) relationship with leaf content of total non-structural carbohydrates (NSC) (A), starch (B), soluble sugar (C), as affected by HLB-disease status in *C. reticulata* ‘W. Murcott’ trees grown under field conditions under protective screen at Stage II of fruit development. Branches were subjected to various girdling and defoliation treatments (not shown due to nonsignificant effects).

### HLB status affected stem response to source-sink manipulation

Stem NSCs and their relationship with leaf and fruit carbohydrates were quantified to investigate their role in shoot carbohydrate dynamics. Stem starch content changed as the fruit developed (*P* < 0.0001; Supplemental Fig. S7B; Supplemental Table S3), with the greatest stem starch content observed at late Stage II. There was also an interaction of source-sink manipulation and HLB disease on stem starch content (*P* = 0.003; Fig. 6). Although, neither healthy nor HLB-affected symptomatic stems accumulated starch in response to girdling. HLB-affected asymptomatic trees’ stem starch content was the most responsive to defoliation treatments, their stems increased starch accumulation when girdled and depleted starch most drastically in response to defoliation.

**Figure 6.**
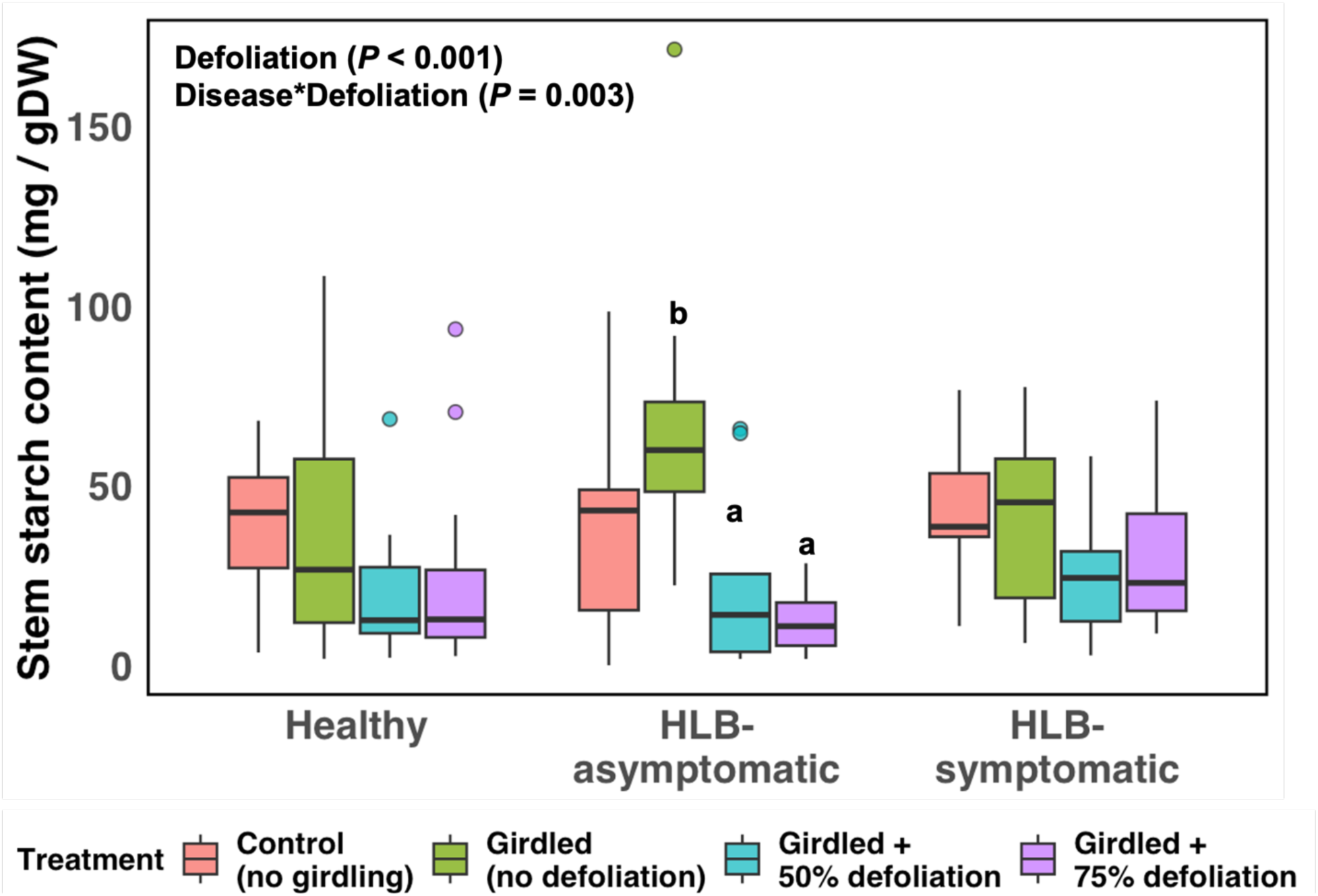
HLB disease and source sink manipulation effect on stem starch content of *C. reticulata* ‘W. Murcott’ trees grown under field conditions under protective screens at Stage II of fruit development. Source-sink manipulations were made on individual branches. Groups sharing different letters differ significantly (Tukey’s honestly significant difference test, *P* < 0.05). To distinguish girdling from defoliation effects, different capital letters indicate significant differences between the ungirdled control and girdled, non-defoliated treatment, whereas different lowercase letters indicate significant differences among defoliation treatments within each disease category.

Stem soluble sugar was significantly affected by HLB disease (*P* = 0.05; Supplemental Fig. S7A), while stem sucrose content was affected by the interaction of HLB disease and source-sink manipulation. Though, stem sucrose content was altered in only HLB-asymptomatic shoots only; specifically stem sucrose reduced in defoliation branches (*P* = 0.001; Supplemental Fig. S8).

We tested correlations of stem NSCs with those of leaves and fruits, followed by linear regression for significant correlations. Stem starch content and soluble sugar were correlated with NSC content in leaves and fruits, as well as with leaf, stem, and fruit dry weights (Fig. 7A-H; Supplemental Fig. S9). We observe a positive relationship between stem starch and leaf or fruit starch, soluble sugar, and dry weights. The exception was fruit starch (Fig. 7B), where increases in fruit starch were associated with reduced stem starch levels. In most cases, HLB shifted the intercept and not the slope of these regressions (7 A, B, F, G and H). The intercept of HLB-symptomatic leaf starch was approximately 50 mg g^-1^ DW higher than those of healthy and HLB-asymptomatic leaves, which were similar. Meanwhile fruit starch and stem soluble sugar intercepts of healthy stems were higher than those of HLB-affected stems. In HLB-asymptomatic trees, fruit and stem dry weight had a higher intercept than both healthy and HLB-symptomatic trees.

**Figure 7:**
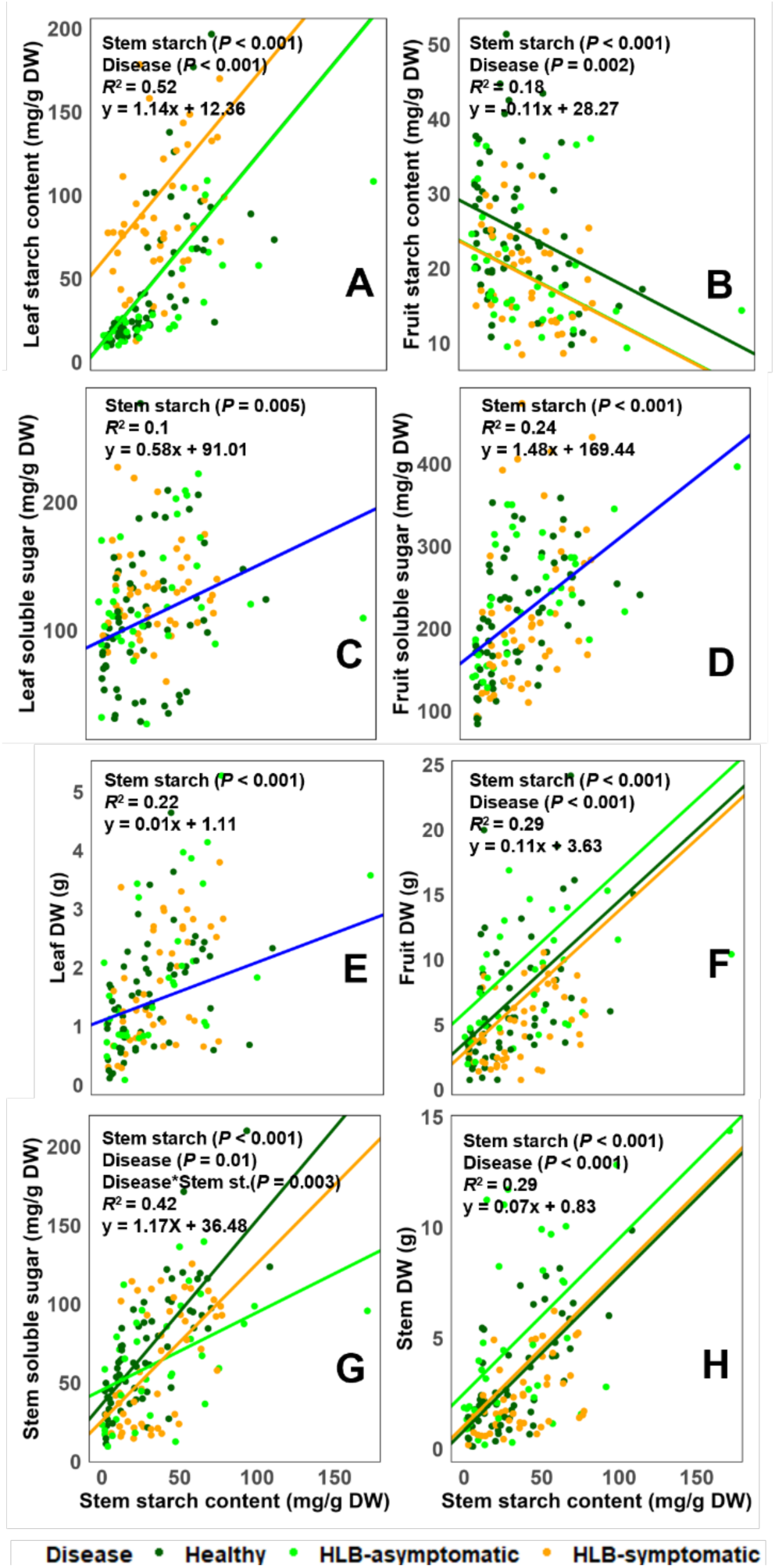
Stem starch relationship with leaf and fruit carbon content as affected by HLB-disease status in *C. reticulata* ‘W. Murcott’ trees grown under protective screens at Stage II of fruit development using a linear mixed effect model at (*P* < 0.05). Linear regression model was fitted, and means were separated with Tukey’s HSD test at P < 0.05. Blue lines represent regression including disease categories. The effects shown are only those that were significant from the model based on analysis of variance. *R*^2^ values refer to the adjusted *R*^2^ for the two-variable relationship shown in the figures

### Changes in the source:sink ratio and HLB influenced fruits independently of each other

To assess whether changes in source:sink manipulation impact fruit carbon dynamics of HLB-affected trees relative to healthy trees, we estimated utilized, accumulated and available carbon by measuring fruit gas exchange, NSCs and fruit dry weight.

Fruit dry weight was affected by sourk-sink manipulation (*P* = 0.04; Fig. 8; Supplemental Table S4) however, no significant differences were detected among the individual treatment means. We observe an interaction effect between HLB and fruit development stage on fruit dry weight (*P* = 0.01; Supplemental Fig. S10). At early Stage II, healthy and HLB-symptomatic fruits had similar dry weights, whereas HLB-asymptomatic fruits had the greatest fruit dry weight. At late Stage II, healthy and HLB-affected asymptomatic fruits had greater fruit dry weight compared to HLB-affected symptomatic fruits (Supplemental Fig. S10).

**Figure 8.**
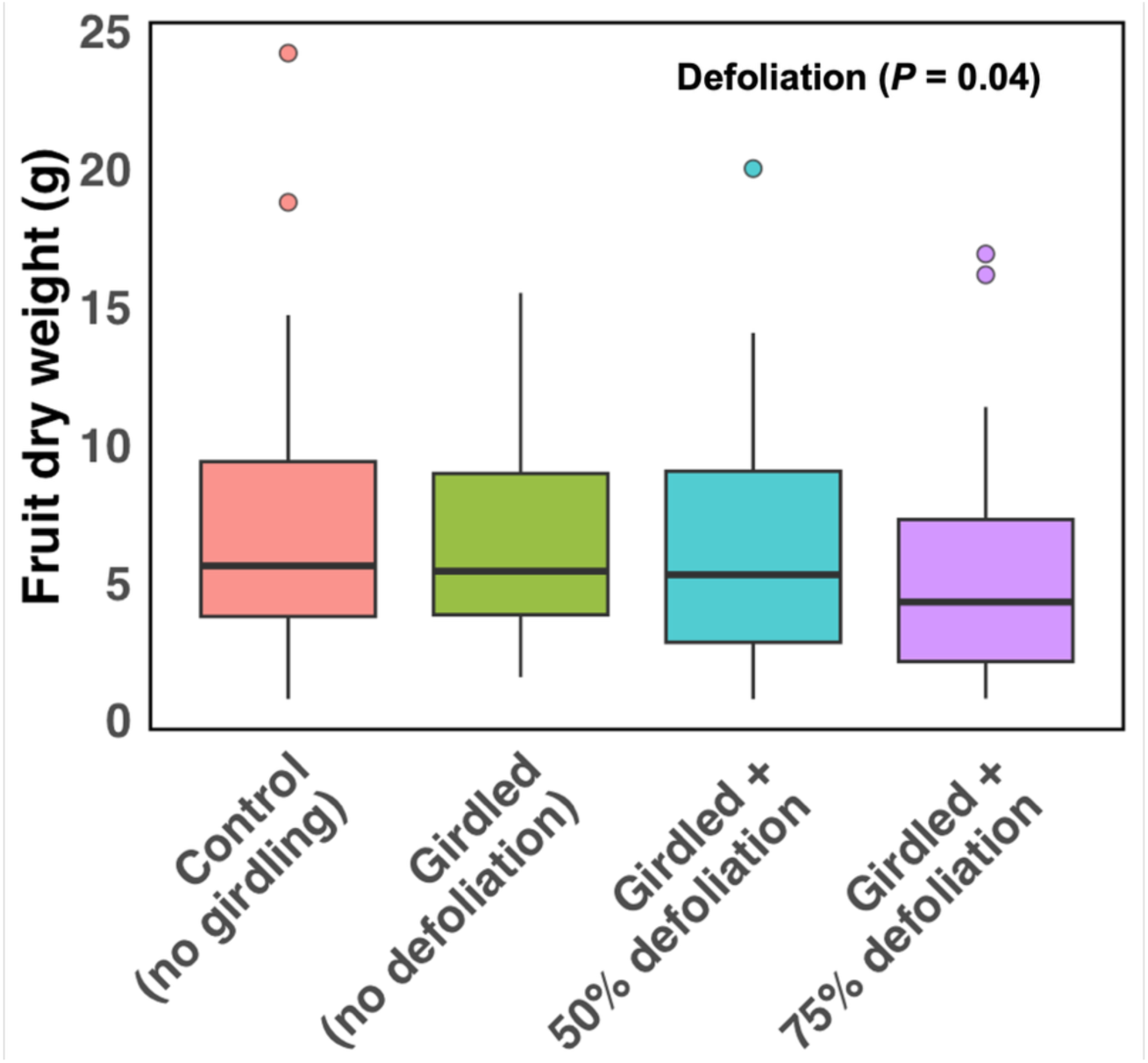
Fruit dry weight of *C. reticulata* ‘W. Murcott’ grown under field condition under protective screens as affected by source-sink manipulation. Source-sink manipulations were made on individual branches.

While fruit sucrose content was not affected by HLB (*P* = 0.3), source-sink manipulation and fruit development stages altered fruit sucrose content (*P* = 0.04 and *P* < 0.001; respectively, Supplemental Fig. S11). Defoliation decreased fruit sucrose level, and it was higher at late stage II of fruit development. Fruit soluble sugar was affected by source-sink manipulation (*P* = 0.007; Supplemental Fig. S12), as girdling increased its content, and defoliation reduced it.

Additionally, HLB reduced fruit starch content (*P* = 0.008; Fig. 9A), and we observed an interaction between source-sink manipulation and fruit developmental stages on fruit starch content (*P*=0.01; Fig. 9B). At early stage II, changes to source:sink ratio affected fruit starch as branch girdling increased fruit starch content, and defoliation reduced it, wheras at late stage II, fruit starch content was similar in all branches irrespective of source-sink manipulation.

**Figure 9.**
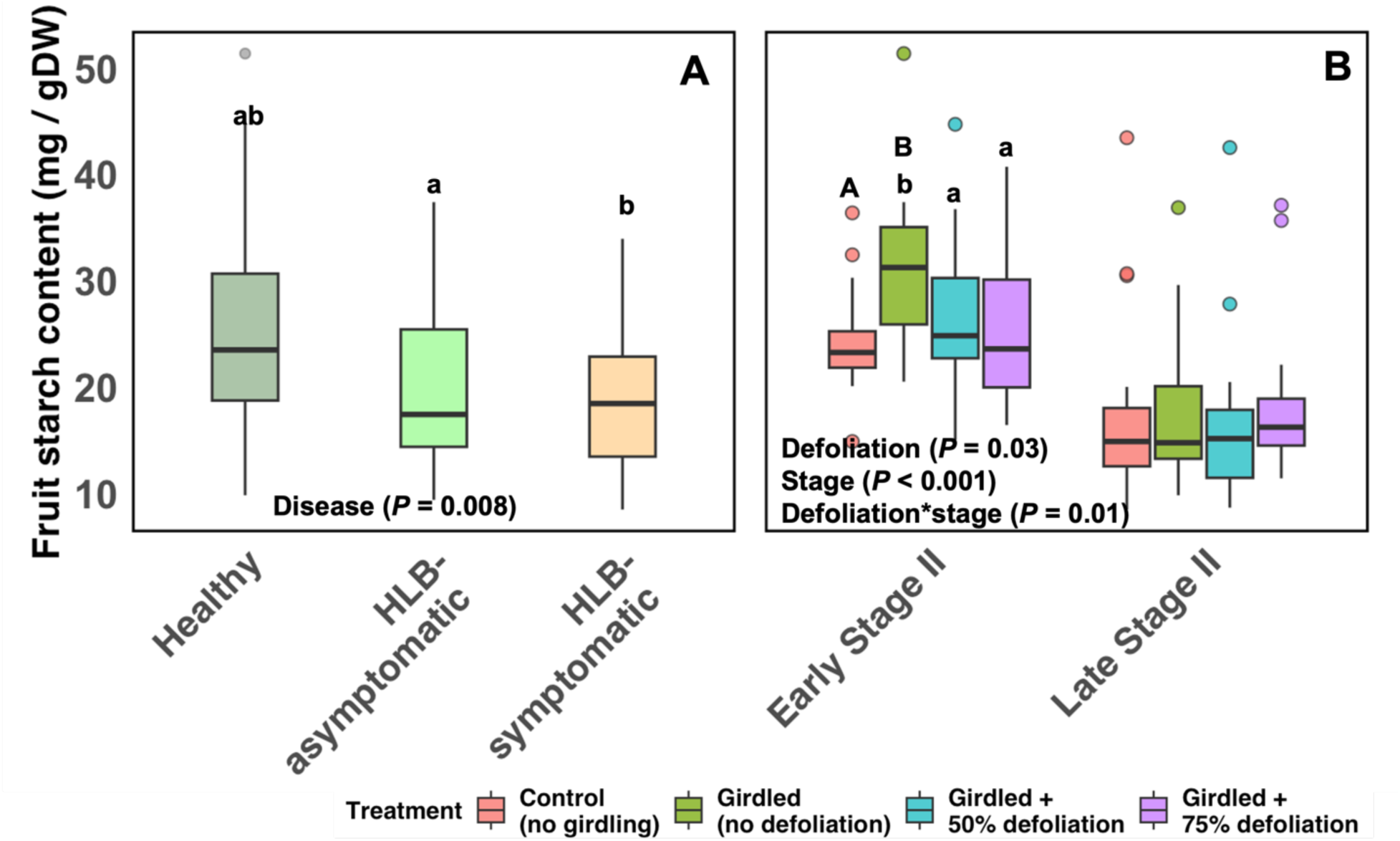
HLB disease (A) and source-sink manipulation across all disease level (B) effect on fruit starch content of *C. reticulata* ‘W. Murcott’ under field condition under protective screens at Stage II of fruit development. Source-sink manipulations were made on individual branches. Groups sharing different letters differ significantly (Tukey’s honestly significant difference test, *P* < 0.05). To distinguish girdling from defoliation effects, different capital letters indicate significant differences between the ungirdled control and girdled, non-defoliated treatment, whereas different lowercase letters indicate significant differences among defoliation treatments within each disease category.

Fruit photosynthesis was impacted only by source-sink manipulation (*P* = 0.005; Fig. 10A) and stages of fruit development (*P* < 0.001; Supplemenatry Fig. S13B), with a marginal effect of HLB on fruit photosynthesis (*P* = 0.06; Supplemental Fig. S13A). Fruit on girdled and ungirdled control branches had similar gross photosynthesis rates, which declined when branches were defoliated, but did not decline firther as defoliation rate increased. Fruit respiration was affected by fruit development stages (*P* < 0.001, Supplemental Fig. S14) and a possible effect of source-sink manipulation (*P* = 0.05; Fig. 10B), and no effect of HLB status on fruit respiration rate.

**Figure 10.**
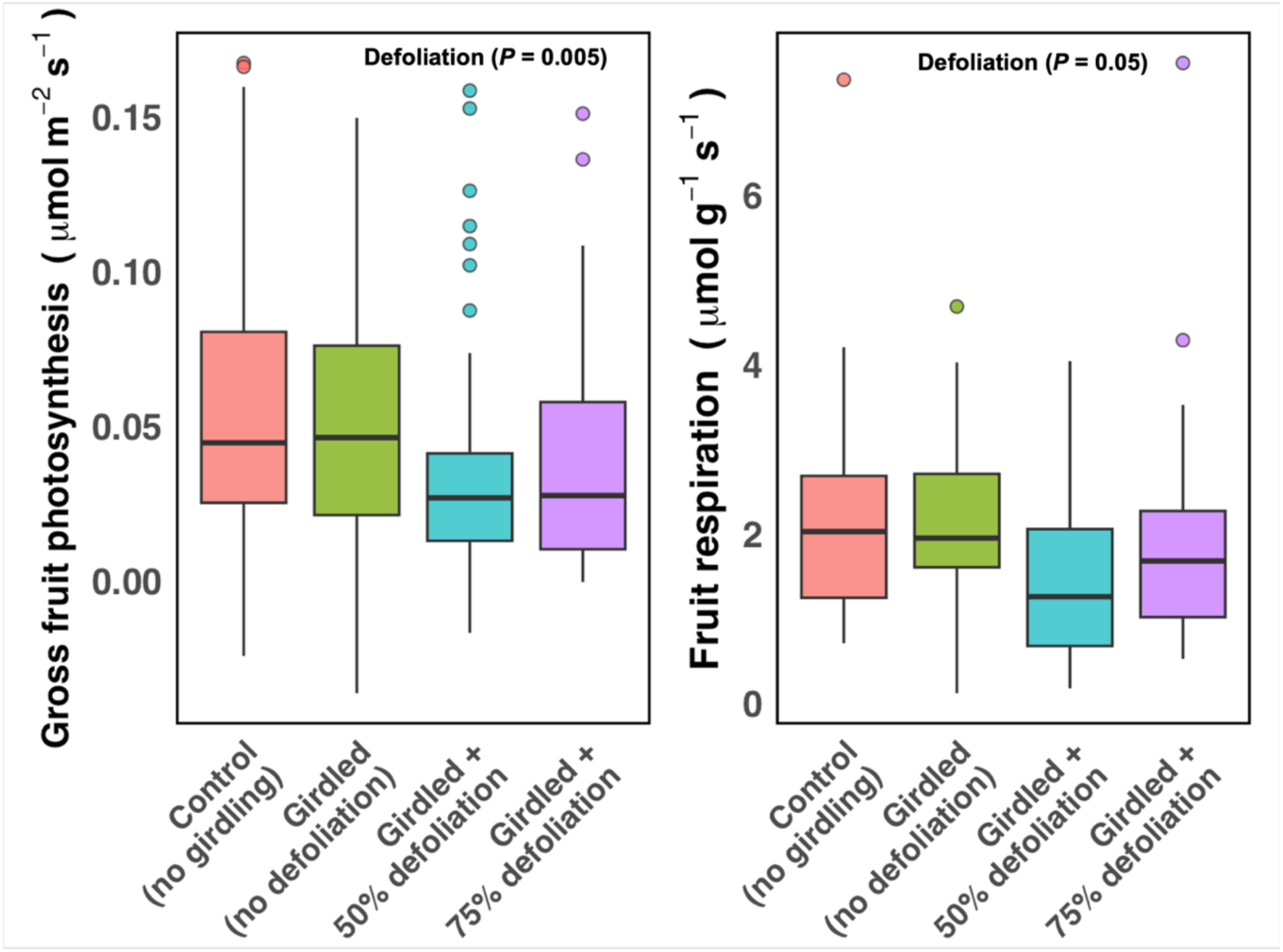
Effect of source-sink manipulation on *C. reticulata* ‘W. Murcott’ fruit photosynthesis per surface area (A) and respiration per fruit dry weight (B) grown on field soil under protective screens across all HLB disease categories. Source-sink manipulations were made on individual branches.

## DISCUSSION

In this study, we investigated the source-sink dynamics of citrus trees and their interaction with HLB. As hypothesized, HLB and source-sink manipulation altered leaf *A*_net_ primarily through stomatal regulation. Stem starch appeared to mediate between leaf and fruit NSCs within and across disease categories. While HLB disease altered fruit growth and carbon accumulation, the shoots’ response to source-sink manipulation were similar across disease categories. Our findings provide a novel perspective on the pattern of HLB disruption of source-sink coordination between leaves and fruits.

### Source-sink photosynthesis response is driven by stomatal regulation

Source-sink regulation of photosynthesis is often mediated by changes in stomatal conductance and biochemical activities (Ainsworth and Bush, 2011; Nebauer *et al*., 2011). As hypothesized, photosynthetic in response to HLB and to manual source-sink manipulation was primarily driven by *g*_s_, and no effect of treatments and disease on Calvin-cycle enzymatic activities was observed. The reduction in *A*_net_ observed in HLB-affected shoots was mostly stomatal, with *R*_dark_ also contributing to HLB-induced reductions in *A*_net_ as additional loss Fig 3). The role of *g*_s_ in source-sink responses is consistent with results of other studies, including whole tree manipulations of citrus (Syvertsen, 1994), and branch girdling in *Coffea arabica* (Avila *et al*., 2020) and in *Prunus avium* (Quentin *et al*., 2013). Keeley *et al*. (2022) posited *g*_s_ as the driver of photosynthetic responses to HLB but did not quantify the relative contributions of Calvin cycle activities, *g*_s_, and *R*_dark_. Contrasting with the present study, studies on whole-tree source-sink manipulation by Syvertsen (1994), found effects of both *g*_s_ and Calvin cycle activities in response to whole tree source-sink manipulations. Turnbull *et al*. (2007) and Li *et al*. (2024) also found strong impacts on Calvin cycle activities rather than *g*_s_. However, studies finding primarily stomatal responses (the present work, Quentin *et al.,* 2013 and Avila *et al*. 2020) involved branch-level manipulations, while those that found Calvin cycle response involved whole-tree manipulation. Thus, a common pattern emerges in which source-sink manipulation at the whole-tree level may induce photosynthetic regulation of Calvin cycle enzymatic activities, while local source-sink responses may involve a greater role of stomatal regulation of photosynthesis. Thus, the spatial scale of source-sink manipulation may influence the specific mechanisms of source-sink attenuation of photosynthesis. Although HLB impacts the entire plant, its photosynthetic response, with *g*_s_ driving reduced *A*_net_, resembles that of branch-level responses rather than whole-plant-level responses, suggesting that leaf-stem coordination may be disrupted together with transport. Further research are required to address what signals or other physiological processes are involved in these scale-dependent responses. If this hypothesis is correct, then HLB would appear to suppress the whole-plant signal for up- or down-regulation of carboxylation, which may explain the partial acclimation of *A* overtime despite increasing disease severity observed by Keeley *et al*. (2022). Symptomatic HLB-infected branches showed distinct photosynthetic responses to source-sink manipulation. *A*_net_ in HLB-symptomatic trees declined sharply between 50% and 75% defoliation, suggesting that acclimation to less-functional transport is not resilient to more extreme source:sink ratio. Thus, HLB photosynthetic responses to source-sink manipulations are driven by *g*_s_, though they may be buffered by the stems as described below.

### Increased leaf R_dark_ and carbohydrate accumulation in relation to HLB symptoms

HLB disease induces callose deposition in the phloem and restricts carbon exports from source tissues. As a result, HLB-affected trees typically show leaf starch accumulation (Welker *et al*., 2022), although the magnitude of this response can vary with disease severity, duration of exposure, or inherent susceptibility (Keeley *et al*., 2022; Robledo *et al.,* 2024). In addition to starch buildup, HLB can also cause subtle but meaningful shifts in leaf soluble sugar and sucrose concentrations (Cai *et al*., 2022), reflecting the disruption of normal source-sink balance.

Although other studies focusing on source-sink manipulation at early stages of fruit development have found source-sink manipulation impacts on foliar sucrose (Iglesias *et al*., 2002; Andrade *et al*., 2019), in this study, despite starch buildup, we did not find any effect of source-sink manipulation on leaf sucrose levels. Thus, the starch pool may have buffered potential fluctuations in sucrose, enabling a sucrose homeostasis, despite HLB infection.

In the present study, leaf carbon content in asymptomatic shoots appear to be more similar to healthy trees than symptomatic trees. Just as in other studies comparing infected asymptomatic and symptomatic plants (Sagaram and Burns, 2009) or leaves (Fan *et al*., 2010), starch accumulated only in the symptomatic leaves. Previous studies reported differences in leaf starch content of healthy and HLB-affected trees based on exposure time (Keeley *et al*., 2022) and tolerance (Robledo *et al.,* 2024). However, in this study, although the trees were exposed to HLB at the same time, starch levels differed between leaves of HLB-asymptomatic and symptomatic trees. Thus, these clear differences between leaves of infected asymptomatic and symptomatic trees may not be associated with disease exposure time or tolerance but to other factors that affect disease progression.

Elevated carbohydrate substrates can stimulate respiration through enhanced metabolic activity (Li *et al*., 2013). In HLB-affected trees, defense response has been linked to elevated maintenance costs such as respiration (Deng *et al*., 2019). Silva *et al*. (2021) reported that HLB increased leaf *R*_dark_ in sweet orange (*C. sinensis* cv. Valencia) trees, a finding our present work confirms and also demonstrating a quantitative relationship between *R*_dark_ and NSC, particularly with soluble sugars. Given that studies have reported that leaf dark respiration is particularly sensitive to concentration of reducing sugars such as glucose, the precursor of pyruvate, which fuels the Krebs cycle (Lee *et al*., 2005; Lu *et al*., 2014). And from the present study, HLB-infected symptomatic leaves had the same slope but a higher intercept of the *R*_dark_ response to soluble sugar as leaves of asymptomatic and uninfected trees. Thus, HLB infection itself may not induce a defense-related increase in respiration. However, symptomatic leaves showed some evidence in support of the hypothesis that HLB induces increase respiration, which is caused by carbohydrate accumulation, thus the increased respiration is associated with symptoms rather than infection. Summarily, HLB affects the homeostasis of foliar sugars, resulting in elevated *R_dark_*, which is independent to source-sink manipulation.

### HLB status affected stem response to source-sink manipulation

The stem plays a dual regulatory role in plant carbon dynamics by functioning as a transient carbon reservoir and a transport conduit. In citrus, the stem has also been identified as an important contributor to total shoot carbon gain due to its capacity for efficient photosynthesis (Keeley *et al*., 2022). In the present study, HLB status affected stem response to source:sink, particularly, HLB-asymptomatic stems showed the strongest adjustments in carbon storage and allocation.

The stem serves as a dynamic buffer managing the ebb and flow of carbon supply and demand between the leaves and fruits (Sala *et al*., 2012; Furze *et al*., 2018). The strong correlations between stem starch and soluble sugar content with the various carbon forms and biomass measured in both fruit and leaves in this study highlights the role of stem in balancing source supply with sink demand. Interestingly, we only observed a negative correlation between stem starch and fruit starch which suggests a competitive relationship between the stem and fruit carbon reserves and likely reflects a trade-off in carbon allocation (Li and Vincent, 2022).

In HLB-affected trees, phloem disruption had previously been reported to increase stem starch accumulation potentially altering how stems respond to changes in the source-sink ratio (Etxeberria *et al*., 2009; Cai *et al*., 2022), although the reports from these studies only showed slight increases in stem starch compared to the hyperaccumulation in leaves. Healthy and HLB-affected trees accumulated similar levels of stem starch, which is reduced by defoliation particularly in HLB-asymptomatic shoots. This finding is comparable with results reported by Robledo *et al*. (2024), who compared sensitive and tolerant genotype responses to HLB, and reported that carbon transport is only slightly reduced in HLB-tolerant cultivars relative to susceptible ones. In summary, these findings suggest that HLB disease alter the stem’s buffering capacity: the most pronounced shifts occur in asymptomatic stems, highlighting the stem’s sensitivity to subtle changes in phloem integrity before visible symptoms appear.

### Changes in the source:sink ratio and disease status influenced fruit traits independently

HLB restricts carbon transport, and reduced carbon delivery to sinks (Johnson *et al*., 2014). As a result, HLB-affected fruits exhibit a range of altered internal and external characteristics, including smaller size, lopsidedness, partially or fully aborted seeds, and an overall reduction in seed number per fruit. These anatomical and morphological changes are accompanied by significant biochemical alterations, such as reduced starch and soluble solids content (Bassanezi *et al*., 2009; Rosales and Burns, 2011; Cai *et al*., 2022; Cifuentes-Arenas *et al*., 2022).

Although HLB alters fruit carbon accumulation and associated physiological traits, fruits of HLB-affected trees did not respond diffrently to source-sink manipulation. The absence of detectable interaction between source-sink manipulation and HLB disease may reflect the relatively short duration of the source-sink manipulation applied in this study relative to the full course of fruit development (8-12 months). Similar to previous studies on differences in fruit dry weight and diameter of healthy and HLB-affected trees, we observed that HLB affected fruit dry weight (Bassanezi *et al*., 2009; Baldwin *et al*., 2010). Symptomatic fruits consistently had the lowest dry weight, likely due to impaired phloem function, which reduces carbon transport to sink tissues. Interestingly, fruits of HLB-asymptomatic trees had similar dry weight as healthy trees despite the presence of *C*Las pathogen, which might be attributed to slower disease progression in HLB-asymptomatic trees and thus less impaired phloem function. Furthermore, fruits of both asymptomatic and symptomatic HLB-affected trees consistently had lower starch content than those of healthy trees. In line with our findings, Cai *et al*. (2022) in their study on HLB impacts on tree physiology of citron (*Citrus medica* L. var. *sarcodactylis* Swingle), also reported that HLB-affected citrus fruits had lower starch content than healthy citron fruits. This is particularly intriguing in fruits of HLB-affected asymptomatic trees, as they maintained similar fruit dry weight to healthy trees despite having less starch. This suggests that HLB-affected trees might allocate carbon to fruit structural carbon over non-structural carbon. Together, these findings indicate that HLB alters fruit carbon accumulation and quality traits, particularly reducing starch content and dry weight in symptomatic fruits. The overall consistency in fruit responses to source-sink manipulation across disease categories suggests that our third hypothesis, that defoliation would more strongly affect HLB-affected fruit than healthy fruit, may be false. HLB appears to affect the overall efficiency of transport which impacts sink activity, while source-sink manipulation further impacts sink activity through total availability of photoassimilates.

## CONCLUSION

The present work presents a source-to-sink assessment of how phloem disease impacts host tree carbon fixation, allocation, and storage showing that: (1) Photosynthetic responses to HLB reflect leaf or branch scale responses and are driven by stomatal regulation. (2) The stem play a central role in carbon distribution, and are the most responsive organ to HLB and to source-sink manipulation, with NSC content correlates with most carbohydrate contents in leaves and fruits. (3) Although HLB alters fruit carbon dynamics, it does not change their response to source-sink manipulation. Summarily, these findings can guide disease and carbon management by identifying key organs for assessing HLB tolerance in citrus, and provides a baseline for comparison with phloem affecting diseases in other species. In the context of this study, the stem is critical, as it is the primary site of HLB interaction with other source-sink dynamics. Moreover, it can inform the development of simple predictive models to estimate how source-sink manipulation and phloem-limited disease alters carbon allocation, organ-specific partitioning for better prediction of plant performance and advancing disease management strategies as well as providing a framework to understanding plant-disease-environment interactions in such diseases.

## Supporting information

Supplementary figures supporting results

Supplementary tables supporting results

## REFERENCES

Achor DS, Etxeberria E, Wang N, Folimonova SY, Chung KR, Albrigo LG. 2010. Sequence of anatomical symptom observations in citrus affected with huanglongbing disease. Plant Pathol J. 9(2):56–64. 10.3923/ppj.2010.56.64

Ainsworth EA, Bush DR. 2011. Carbohydrate export from the leaf: a highly regulated process and target to enhance photosynthesis and productivity. Plant Physiol. 155(1):64–69. 10.1104/pp.110.167684

Andrade D, Covarrubias MS, Benedetto G, Moya CB, Troncoso-Ponce B. 2019. Differential source–sink manipulation affects leaf carbohydrate and photosynthesis of early- and late-harvest nectarine varieties. Theor Exp Plant Physiol. 31(3):341–356. 10.1007/s40626-019-00150-0

Avila RT, Martins SCV, Sanglard LMVP, dos Santos MS, Menezes-Silva PE, Detman KC, Sanglard ML, Cardoso AA, Morais LE, Vital CE, et al. 2020. Starch accumulation does not lead to feedback photosynthetic downregulation in girdled coffee branches under varying source-to-sink ratios. Trees. 34:1–16. 10.1007/s00468-019-01893-8

Baldwin E, Plotto A, Manthey J, McCollum G, Bai J, Irey M, Cameron R, Luzio G. 2010. Postharvest treatments for huanglongbing (HLB)-affected citrus fruit to reduce peel greening and improve fresh market appearance. J Agric Food Chem. 58(2):1247–1262. 10.1021/jf9031958

Bassanezi RB, Montesino LH, Stuchi ES. 2009. Effects of huanglongbing on fruit quality of sweet orange cultivars in Brazil. Eur J Plant Pathol. 125(4):565–572. 10.1007/s10658-009-9506-3

Bates D, Mächler M, Bolker B, Walker S. 2015. Fitting linear mixed-effects models using lme4. J Stat Softw. 67(1):1–48. 10.18637/jss.v067.i01

Bendix C, Lewis JD. 2018. The enemy within: phloem-limited pathogens. Mol Plant Pathol. 19:238–254. 10.1111/mpp.12526

Cai Q, Liu C, Yuan M, Sun L, Yang Q, Zhou L. 2022. HLB induce changes in the tree physiology of citron (Citrus medica L. var. sarcodactylis Swingle). Physiol Mol Biol Plants. 28(2):517–531. 10.1007/s12298-022-01129-z

Cifuentes-Arenas JC, de Oliveira HT, Raiol-Júnior LL, de Carvalho EV, Kharfan D, Creste AL, Gastaminza G, Salas H, Bassanezi RB, Ayres AJ, et al. 2022. Impacts of huanglongbing on fruit yield and quality and on flushing dynamics of Sicilian lemon trees. Front Plant Sci. 13:1005557. 10.3389/fpls.2022.1005557

Deng H, Achor D, Etxeberria E, Yu Q, Du D, Stanton D, Liang G, Gmitter FG Jr. 2019. Phloem regeneration is a mechanism for Huanglongbing-tolerance of ‘Bearss’ lemon and ‘LB8-9’ Sugar Belle® mandarin. Front Plant Sci. 10:277. 10.3389/fpls.2019.00277

Etxeberria E, Gonzalez P, Achor D, Albrigo G. 2009. Anatomical distribution of abnormally high levels of starch in HLB-affected Valencia orange trees. Physiol Mol Plant Pathol. 74(1):76–83. 10.1016/j.pmpp.2009.09.004

Fan J, Chen C, Brlansky RH, Gmitter FG Jr, Li ZG. 2010. Changes in carbohydrate metabolism in Citrus sinensis infected with ‘Candidatus Liberibacter asiaticus’. Plant Pathol. 59:1037–1043. 10.1111/j.1365-3059.2010.02328.x

Furze ME, Trumbore S, Hartmann H. 2018. Detours on the phloem sugar highway: stem carbon storage and remobilization. Curr Opin Plant Biol. 43:89–95. 10.1016/j.pbi.2018.02.005

Gessler A, Zweifel R. 2024. Beyond source and sink control—toward an integrated approach to understand the carbon balance in plants. New Phytol. 242(2):858–869. 10.1111/nph.19611

Graham J, Gottwald T, Setamou M. 2020. Status of Huanglongbing (HLB) outbreaks in Florida, California and Texas. Trop Plant Pathol. 45(3):265–278. 10.1007/s40858-020-00335-y

Gross J, Gallinger J, Görg LM. 2022. Interactions between phloem-restricted bacterial plant pathogens, their vector insects, host plants, and natural enemies, mediated by primary and secondary plant metabolites. Entomol Gen. 42(2):185–215. 10.1127/entomologia/2021/1254

Hothorn T, Bretz F, Westfall P. 2008. Simultaneous inference in general parametric models. Biom J. 50(3):346–363. 10.1002/bimj.200810425

Iglesias DJ, Lliso I, Tadeo FR, Talon M. 2002. Regulation of photosynthesis through source imbalance in citrus is mediated by carbohydrate content in leaves. Physiol Plant. 116(4):563–572. 10.1034/j.1399-3054.2002.1160416.x

Inoue H, Yamashita S, Muraki K, Fujiwara K, Honda H, Ono T, Nonaka Y, Kato T, Matsuyama S, Sugano M, et al. 2020. Fe2+ ions alleviate the symptom of citrus greening disease. Int J Mol Sci. 21(11):4033. 10.3390/ijms21114033

Johnson EG, Wu J, Bright DB, Graham JH. 2014. Association of ‘Candidatus Liberibacter asiaticus’ root infection, but not phloem plugging, with root loss on huanglongbing-affected trees prior to appearance of foliar symptoms. Plant Pathol. 63:290–298. 10.1111/ppa.12109

Kaur A, Zhang L, Yang M, Maness N, Graham CJ, Kumari R, Sun Y, Panta S, Ferguson L. 2023. Evaluation of natural spring freeze tolerance of five pecan scion–rootstock combinations. HortScience. 58(10):1139–1148. 10.21273/HORTSCI17258-23

Keeley M, Rowland D, Vincent C. 2022. Citrus photosynthesis and morphology acclimate to phloem-affecting huanglongbing disease at the leaf and shoot levels. Physiol Plant. 174(2). 10.1111/ppl.13662

Khalilzadeh M, Kumari R, Alade DO, Levy A, Vincent C. 2026. Fitting the pieces into a whole picture: physiology and pathogenesis in huanglongbing disease. Physiol Mol Plant Pathol. 146:103476. 10.1016/j.pmpp.2026.103476

Kunta M, da Graça JV, Malik NS, Louzada ES, Sétamou M. 2014. Quantitative distribution of ‘Candidatus Liberibacter asiaticus’ in the aerial parts of the huanglongbing-infected citrus trees in Texas. HortScience. 49(1):65–68. 10.21273/HORTSCI.49.1.65

Lee TD, Reich SB, Bolstad SV. 2005. Acclimation of leaf respiration to temperature is rapid and related to specific leaf area, soluble sugars and leaf nitrogen across three temperate deciduous tree species. Funct Ecol. 19(4):640–647. 10.1111/j.1365-2435.2005.01025.x

Lemoine R, La Camera S, Atanassova R, Dédaldéchamps F, Allario T, Pourtau N, Bonnemain JL, Laloi M, Coutos-Thévenot P, Maurousset L, et al. 2013. Source-to-sink transport of sugar and regulation by environmental factors. Front Plant Sci. 4:272. 10.3389/fpls.2013.00272

Levy A, Livingston T, Wang C, Achor D, Vashisth T. 2023. Canopy density, but not bacterial titers, predicts fruit yield in huanglongbing-affected sweet orange trees. Plants. 12(2):290. 10.3390/plants12020290

Li S, Hussain SB, Vincent C. 2024. Response of carbon fixation, allocation, and growth to source-sink manipulation by defoliation in vegetative citrus trees. Physiol Plant. 176(3). 10.1111/ppl.14304

Li S, Vincent C. 2022. Root balance controls flush phenology and carbohydrate translocation dynamics in citrus (Citrus × sinensis) trunk. Physiol Plant. 174. 10.1111/ppl.13601

Li T, Heuvelink E, Marcelis LFM. 2015. Quantifying the source-sink balance and carbohydrate content in three tomato cultivars. Front Plant Sci. 6:416. 10.3389/fpls.2015.00416

Li X, Zhang G, Sun B, Zhang S, Zhang Y, Liao Y, Zhou Y, Xia X, Shi K, Yu J. 2013. Stimulated leaf dark respiration in tomato in an elevated carbon dioxide atmosphere. Sci Rep. 3:3433. 10.1038/srep03433

Lu YB, Yang LT, Li Y, Xu J, Liao TT, Chen YB, Chen LS. 2014. Effects of boron deficiency on major metabolites, key enzymes and gas exchange in leaves and roots of Citrus sinensis seedlings. Tree Physiol. 34(6):608–618. 10.1093/treephys/tsu047

Matias P, Barrote I, Azinheira G, Continella A, Duarte A. 2023. Citrus pruning in the Mediterranean climate: a review. Plants. 12(19):3360. 10.3390/plants12193360

Nebauer SG, Renau-Morata B, Guardiola JL, Molina RV. 2011. Photosynthesis down-regulation precedes carbohydrate accumulation under sink limitation in citrus. Tree Physiol. 31(2):169–177. 10.1093/treephys/tpq103

Paul MJ, Foyer CH. 2001. Sink regulation of photosynthesis. J Exp Bot. 52(360):1383–1400. 10.1093/jexbot/52.360.1383

Quentin AG, Close DC, Hennen LMHS, Pinkard EA. 2013. Down-regulation of photosynthesis following girdling, but contrasting effects on fruit set and retention, in two sweet cherry cultivars. Plant Physiol Biochem. 73:359–367. 10.1016/j.plaphy.2013.10.014

Robledo J, Welker S, Shtein I, Bernardini C, Vincent C, Levy A. 2024. Phloem and xylem responses are both implicated in HLB tolerance of Sugar Belle. Phytopathology. 114(2): 441–453. 10.1094/PHYTO-05-23-0148-R

Rosales R, Burns JK. 2011. Phytohormone changes and carbohydrate status in sweet orange fruit from huanglongbing-infected trees. J Plant Growth Regul. 30(3):312–321. 10.1007/s00344-011-9193-0

Sagaram M, Burns JK. 2009. Leaf chlorophyll fluorescence parameters and huanglongbing. J Am Soc Hortic Sci. 134(2):194–201. 10.21273/JASHS.134.2.194

Sala A, Woodruff DR, Meinzer FC. 2012. Carbon dynamics in trees: feast or famine? Tree Physiol. 32(6):764–775. 10.1093/treephys/tpr143

Schloerke B, Cook D, Larmarange J, Briatte J, Marbach M, Thoen E, Elberg A, Crowley J. 2025. GGally: extension to ‘ggplot2’ [R package]. Version 2.4.0. Available from: https://ggobi.github.io/ggally/

Silva JRD, Boaretto RM, Lavorenti JAL, dos Santos BCF, Coletta-Filho HD, Mattos D. 2021. Effects of deficit irrigation and huanglongbing on sweet orange trees. Front Plant Sci. 12:731314. 10.3389/fpls.2021.731314

Singerman A, Rogers ME. 2020. The economic challenges of dealing with citrus greening: the case of Florida. J Integr Pest Manag. 11(1):3. 10.1093/jipm/pmz037

Stinziano JR, McDermitt DK, Lynch DJ, Saathoff AJ, Morgan PB, Hanson DT. 2019. The rapid A/Ci response: a guide to best practices. New Phytol. 221:625–627. 10.1111/nph.15448

Syvertsen JS. 1994. Partial shoot removal increases net CO2 assimilation and alters water relations of citrus seedlings. Tree Physiol. 14(5):497–508. 10.1093/treephys/14.5.497

Tang L, Singh S, Vashisth T. 2020. Association between fruit development and mature fruit drop in huanglongbing-affected sweet orange. HortScience. 55(6):851–857. 10.21273/HORTSCI14931-20

Tipu MMH, Masud MM, Jahan R, Baroi A, Hoque AKMA. 2021. Identification of citrus greening based on visual symptoms: a grower’s diagnostic toolkit. Heliyon. 7(11). 10.1016/j.heliyon.2021.e08387

Turnbull TL, Adams MA, Warren CR. 2007. Increased photosynthesis following partial defoliation of field-grown Eucalyptus globulus seedlings is not caused by increased leaf nitrogen. Tree Physiol. 27(10):1481–1492. 10.1093/treephys/27.10.1481

Welker S, Pierre M, Santiago JP, Dutt M, Vincent C, Levy A. 2022. Phloem transport limitation in huanglongbing-affected sweet orange is dependent on phloem-limited bacteria and callose. Tree Physiol. 42(2):379–390. 10.1093/treephys/tpab134

White AC, Rogers A, Rees M, Osborne CP. 2016. How can we make plants grow faster? A source-sink perspective on growth rate. J Exp Bot. 67(1):31–45. 10.1093/jxb/erv447

Zhao M, Liu Z, Hu Y, Yi S, Zhang Y, Hu B, Shi X, Rennenberg H. 2025. Carbon metabolism and partitioning in citrus leaves is determined by hybrid, cultivar and leaf type. Plant Physiol Biochem. 224:109978. 10.1016/j.plaphy.2025.109978

