## Supplementary figures supporting results for "Phloem-Limited Disease Impacts Citrus Carbohydrate Dynamics Distinctly from Source-Sink Manipulation"

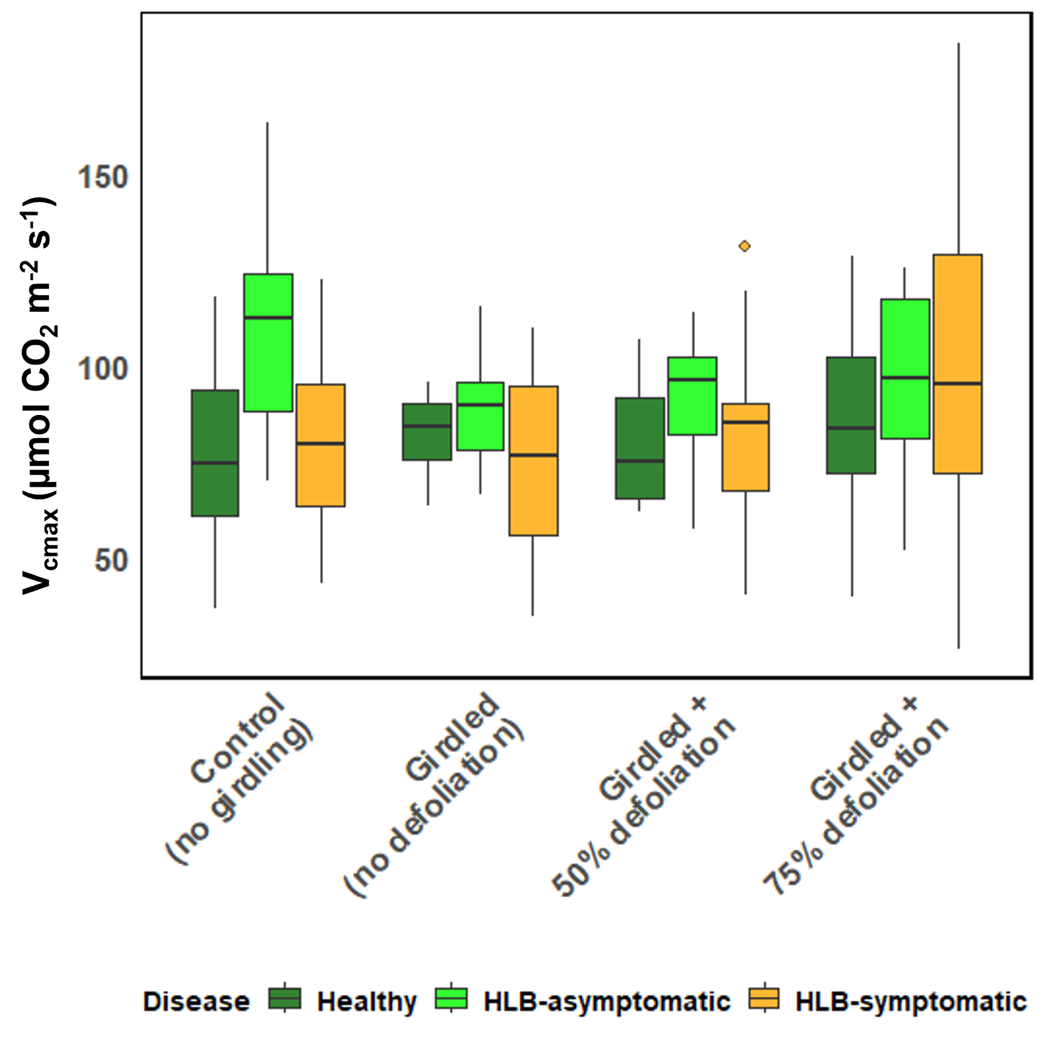

Figure S1: Effect of HLB disease and source-sink manipulation on *V_c_*_max_ (maximum carboxylation rate of rubisco) in leaves of field grown *C. reticulata* ‘W. Murcott’ under protective screens at Stage II of fruit development. The disease status of the trees was determined using qPCR assays and source-sink manipulations were made on individual branches. Each point represents the mean of *V_c_*_max_, the lines showing the trends across defoliation and diseases and the error bar shows the standard error (n=6).

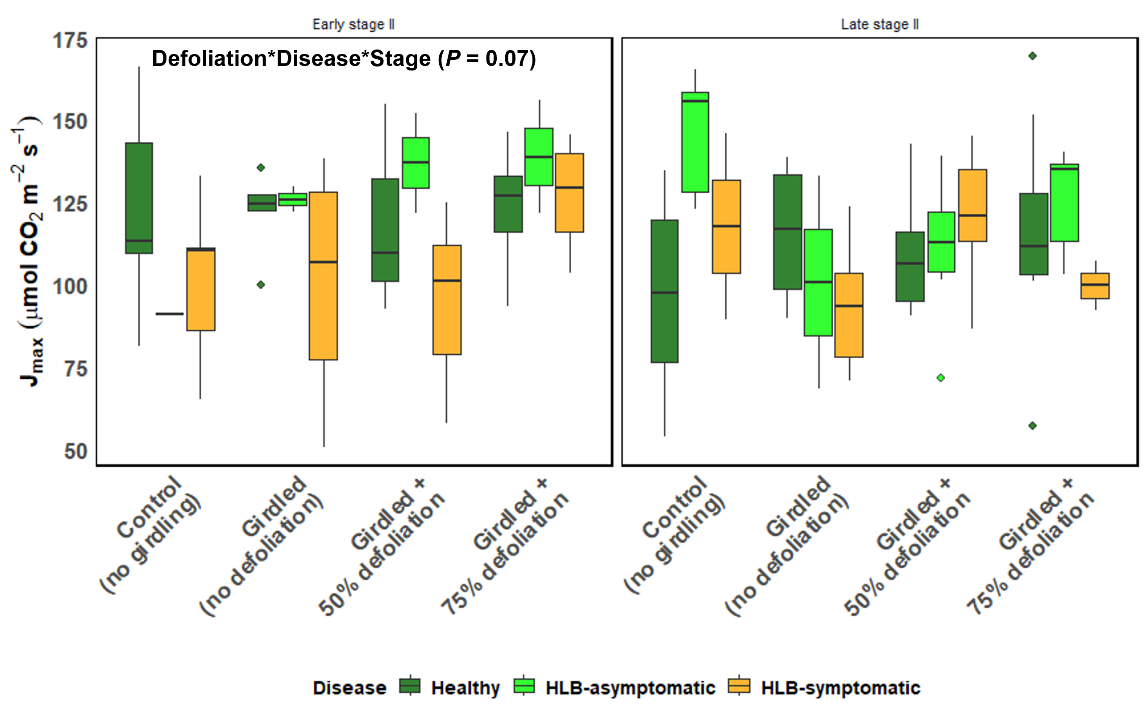

Figure S2**:** Effect of HLB disease and source-sink manipulation at early and late Stage II of fruit development on *J*_max_ (photosynthetic electron transport rate) in leaves of field grown *C. reticulata* ‘W. Murcott’ grown under protective screens. The disease status of the trees was determined using qPCR assays and source-sink manipulations were made on individual branches.

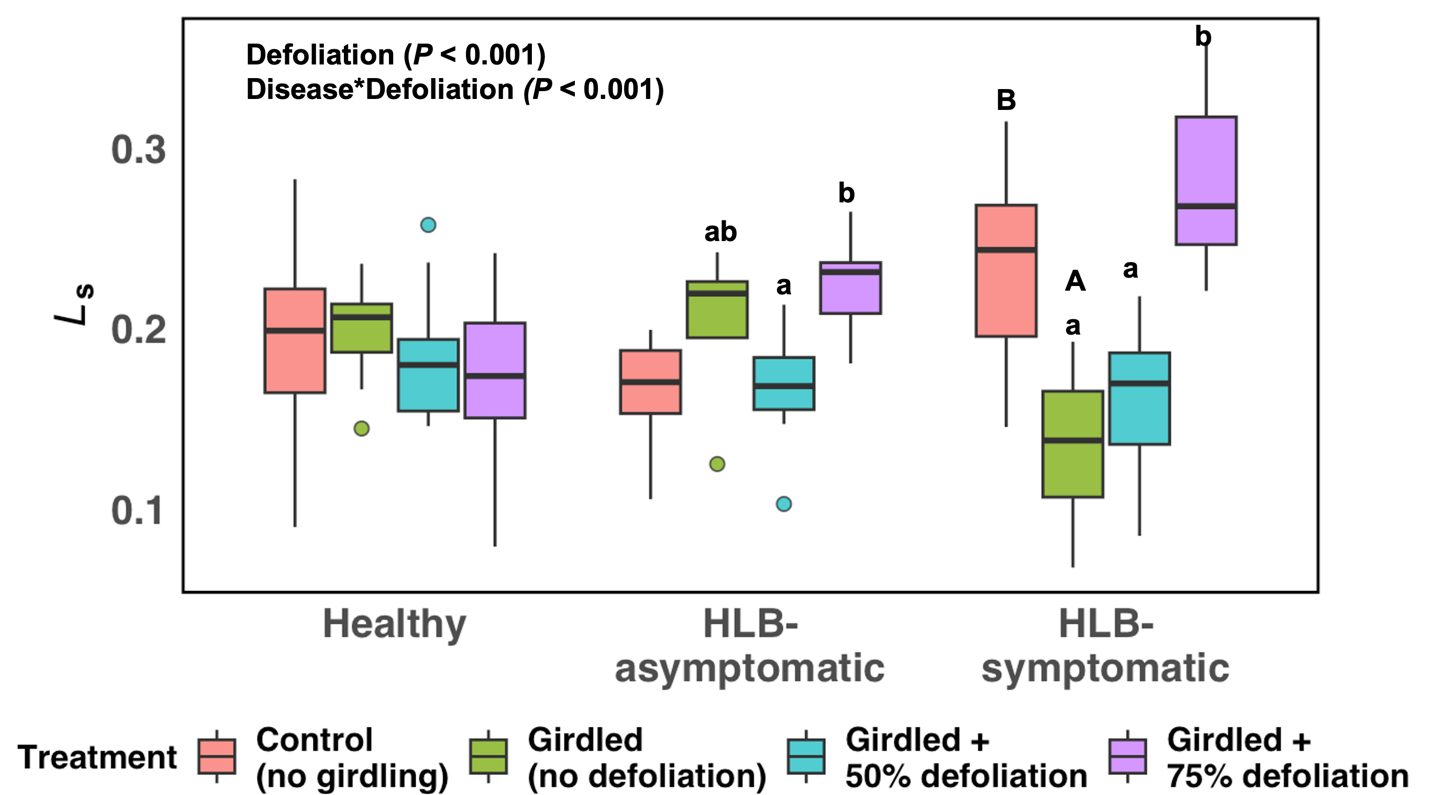

Figure S3**:** Stomatal limitation (*L*_s_) at early and late Stage II of fruit development of field grown *C. reticulata* ‘W. Murcott’ trees grown under protective screens as affected by HLB disease and source-sink manipulation. The disease status of the trees was determined using qPCR assays. Source-sink manipulations were made on individual branches. Groups sharing different letters differ significantly (Tukey’s honestly significant difference test, *P* < 0.05). To distinguish girdling from defoliation effects, different capital letters indicate significant differences between the ungirdled control and girdled, non-defoliated treatment, whereas different lowercase letters indicate significant differences among defoliation treatments within each disease category.

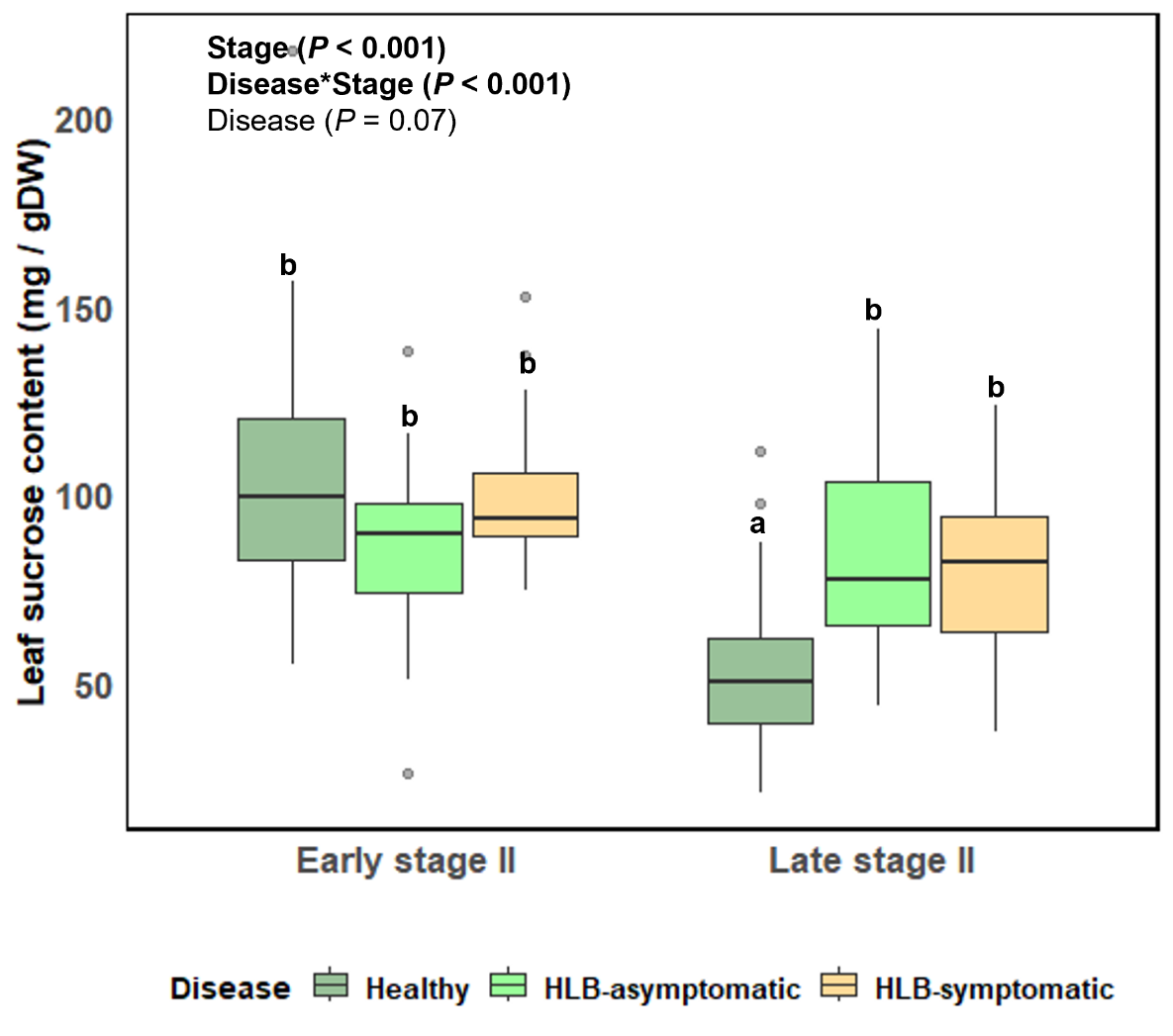

Figure S4**:** Leaf sucrose content as impacted by HLB disease at early and late Stage II of fruit development in *C. reticulata* ‘W. Murcott’ trees grown under field conditions under protective screens. Groups labeled with different letters are significantly different (Tukey’s Honestly Significant Difference; *P* < 0.05).

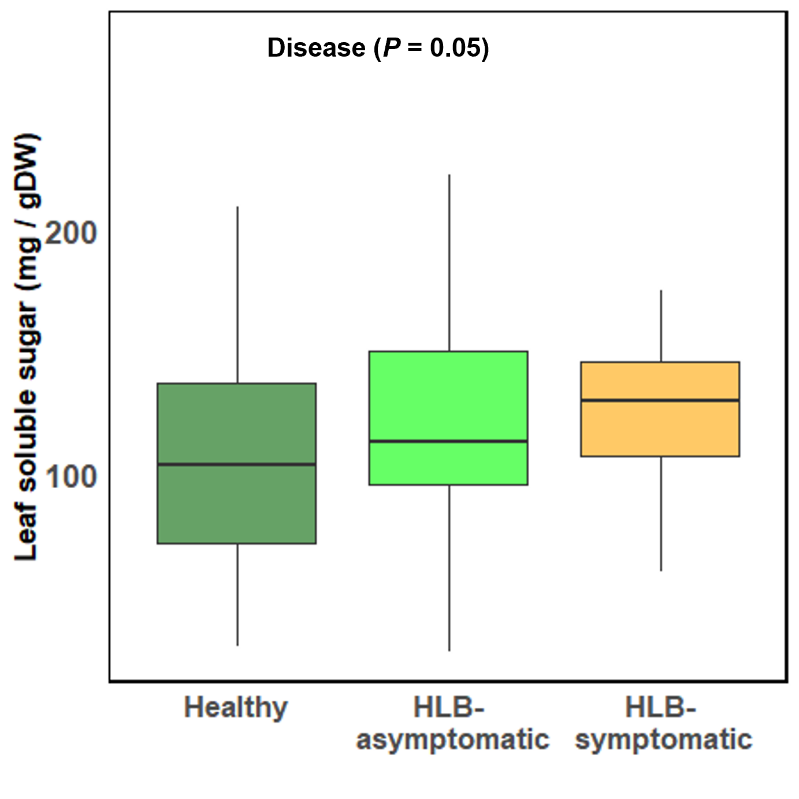

Figure S5**:** Effect of HLB disease on leaf soluble sugar at early and late Stage II of fruit development of *C. reticulata* ‘W. Murcott’ trees grown under field conditions under protective screens at varying source:sink ratio.

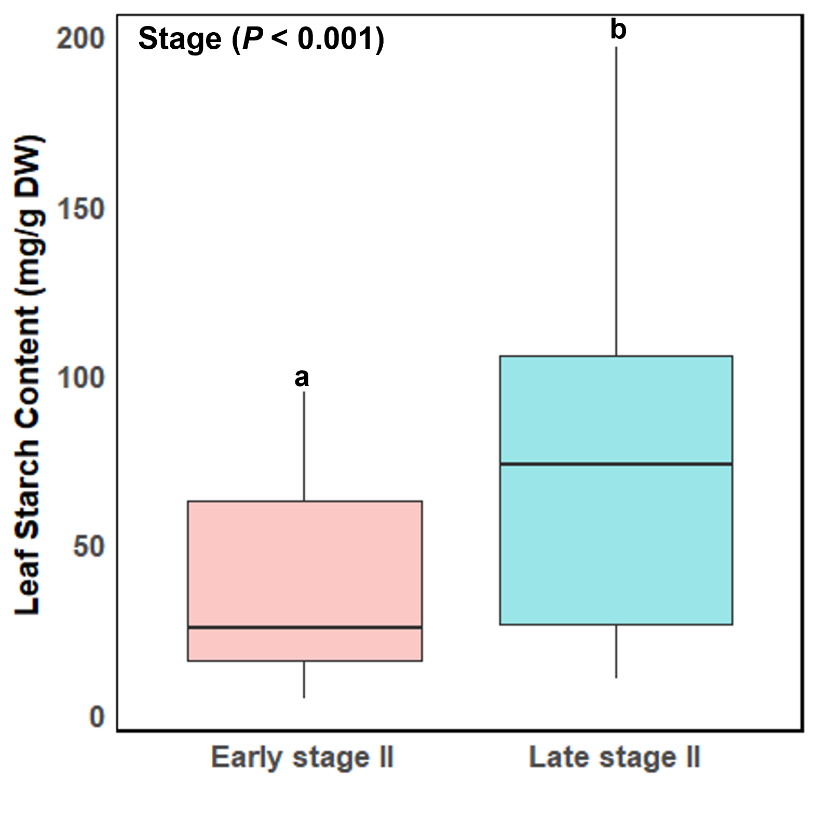

Figure S6**:** Leaf starch content of healthy and HLB diseased *C. reticulata* ‘W. Murcott’ trees grown field under field conditions under protective screens at early and late Stage II of fruit development with varying source-sink manipulated. Groups labeled with different letters are significantly different (Tukey’s Honestly Significant Difference; *P* < 0.05).

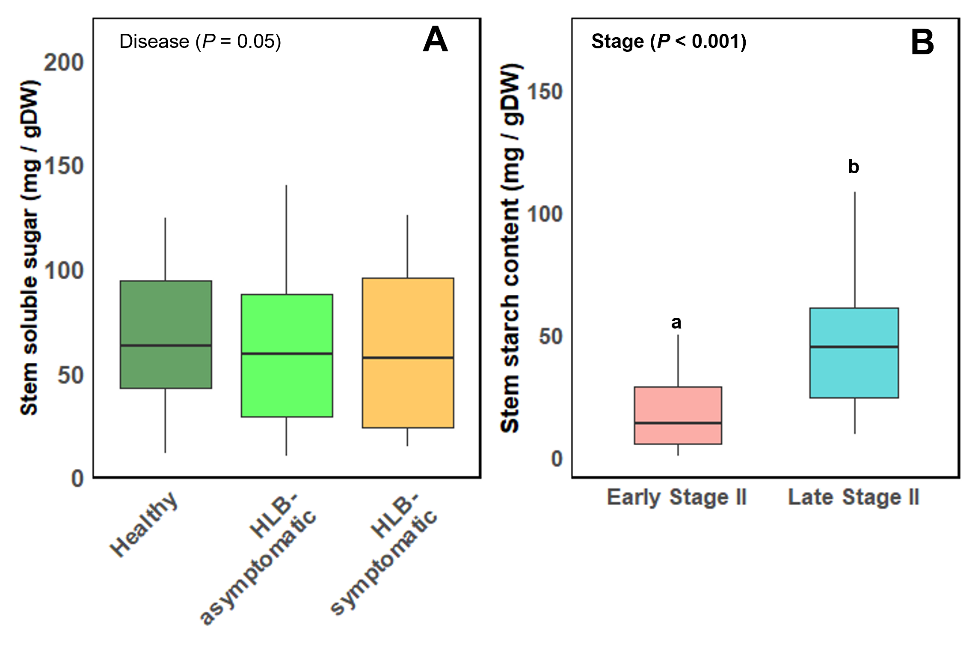

Figure S7: Stem soluble sugar (A) starch content (B) at early and late Stage II of fruit development of healthy and HLB diseased of field grown *C. reticulata* ‘W. Murcott’ trees grown under protective screens at varying source:sink manipulation. Groups labeled with different letters are significantly different (Tukey’s Honestly Significant Difference; *P* < 0.05).

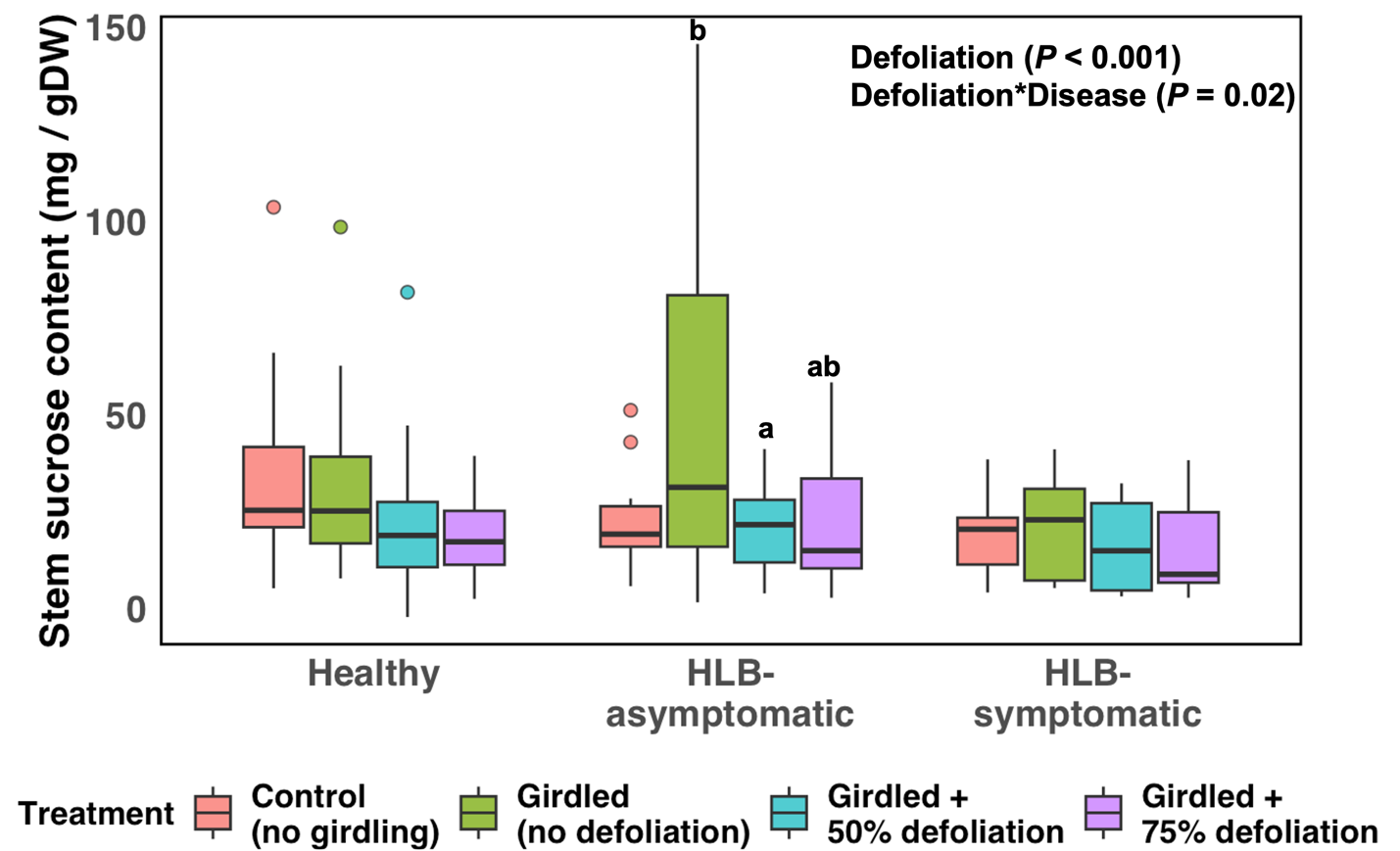

Figure S8: Effect of source-sink manipulation and HLB disease on stem sucrose content in healthy and HLB diseased of field grown *C. reticulata* ‘W. Murcott’ grown under protective screens. Source-sink manipulations were made on individual branches. Groups sharing different letters differ significantly (Tukey’s honestly significant difference test, *P* < 0.05). To distinguish girdling from defoliation effects, different capital letters indicate significant differences between the ungirdled control and girdled, non-defoliated treatment, whereas different lowercase letters indicate significant differences among defoliation treatments within each disease category.

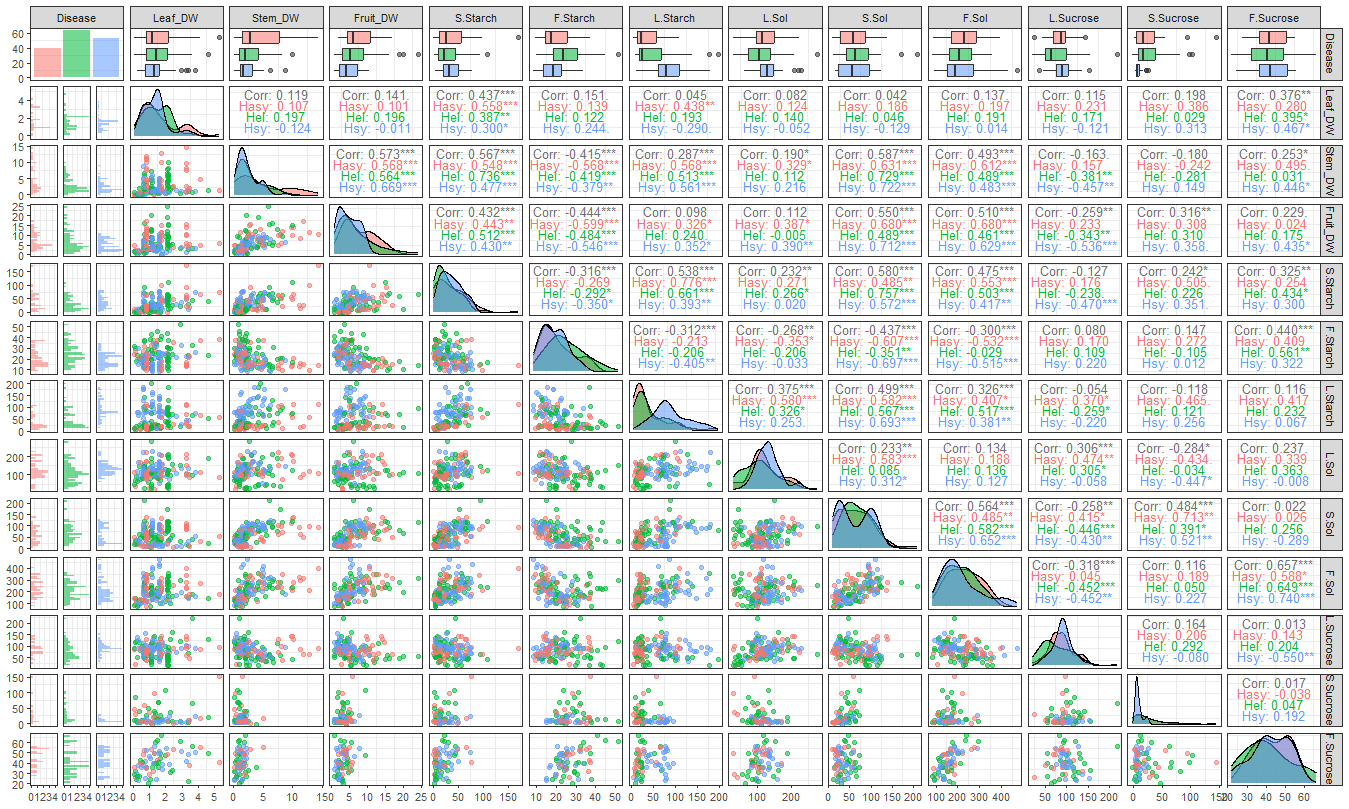

Figure S9: Non-structural carbon and dry weight correlations of leaves, stems, and fruit of field grown *C. reticulata* ‘W. Murcott’ healthy and HLB-affected trees grown under protective screens at varying source:sink ratio. DW- dry weight in g; S – Stem; F- Fruit; L- Leaf; Sol- Soluble sugar, Hel- Healthy (green color); Hasy- HLB-asymptomatic (red color) and Hsy- HLB-symptomatic (blue color).

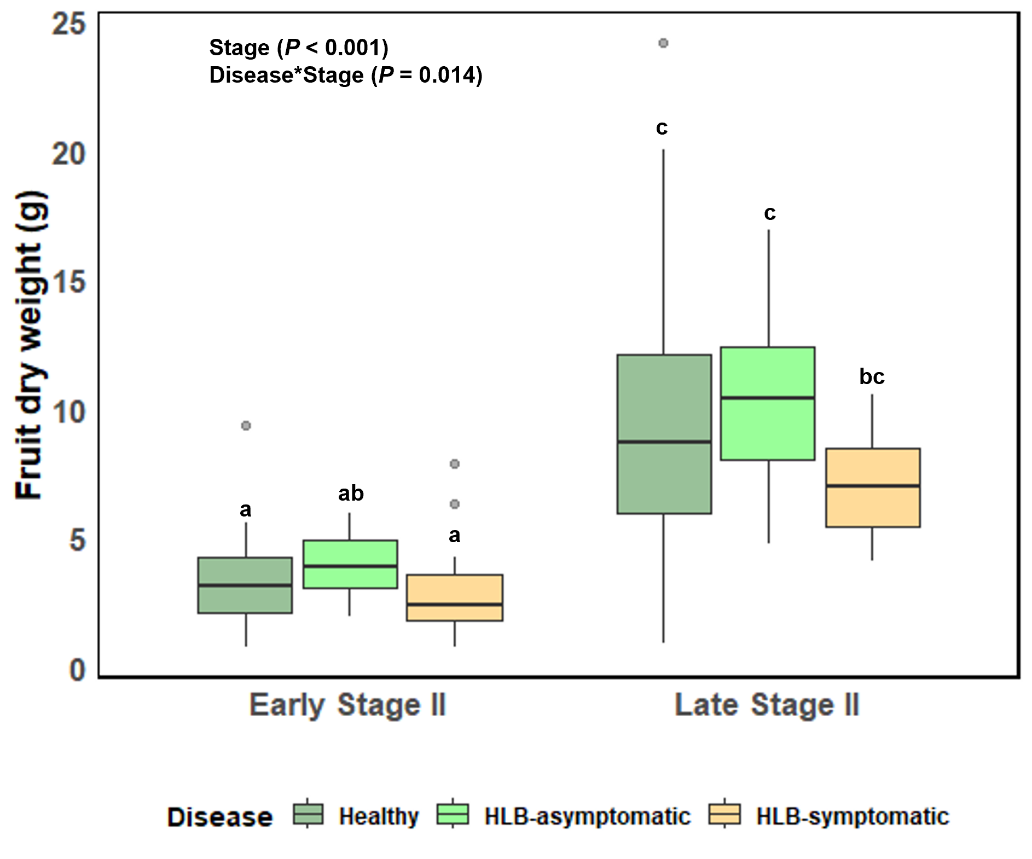

Figure S10**:** Fruit dry weight of field grown *C. reticulata* ‘W. Murcott’ trees grown under protective screens at early and late Stage II of fruit development. Groups with different letters at significantly different (Tukey’s Honestly Significant Difference; *P* < 0.05).

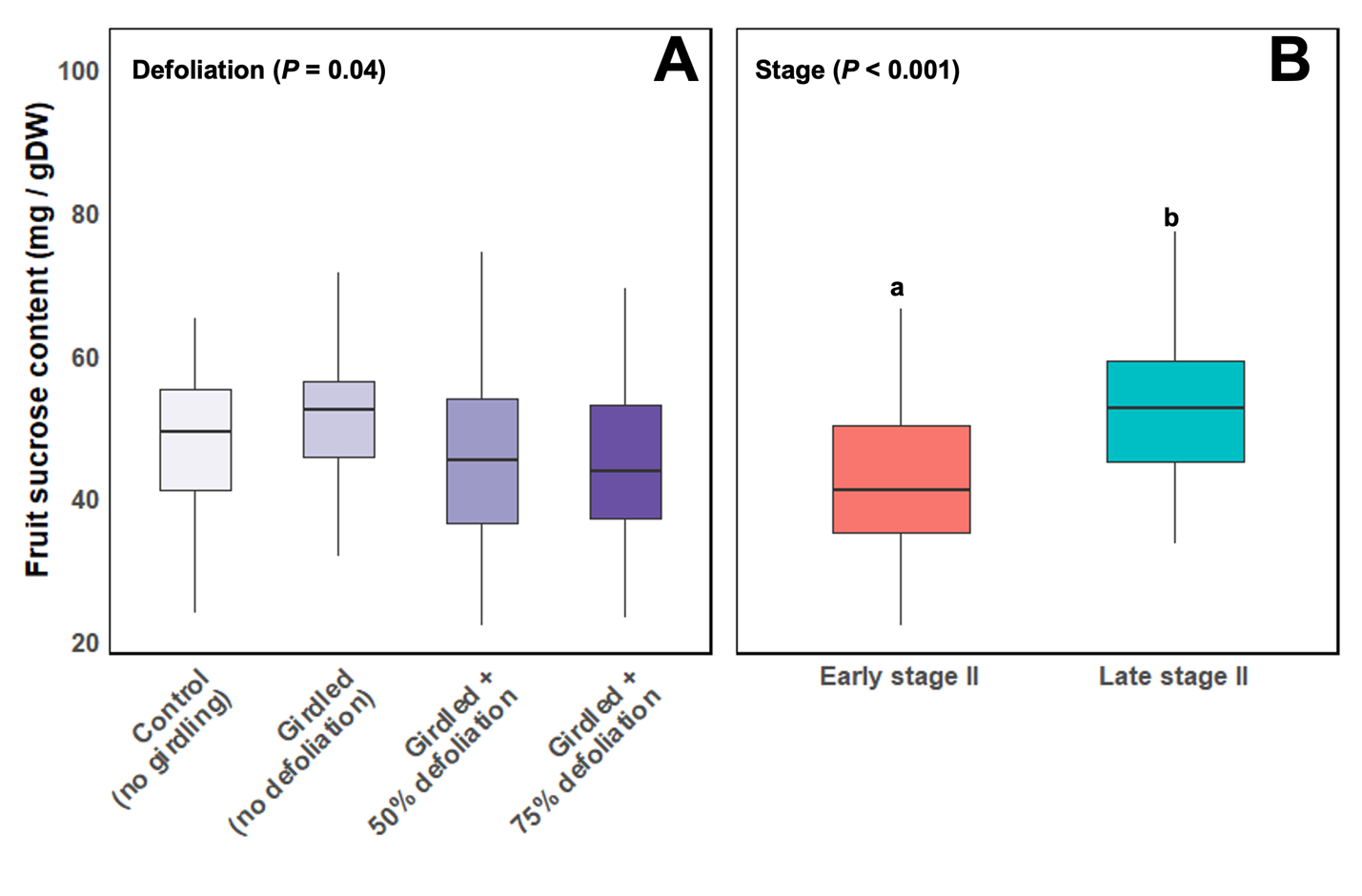

Figure S11**:** Effect of source-sink manipulation (A) at early and late Stage II of fruit development (B) on fruit sucrose content of field grown *C. reticulata* ‘W. Murcott’ trees grown under field conditions under protective screens. Source-sink manipulations were made on individual branches, analyzed using Tukey’s honestly significant difference at *P* < 0.05. Groups sharing different letters differ significantly (Tukey’s honestly significant difference test, *P* < 0.05). To distinguish girdling from defoliation effects, different capital letters indicate significant differences between the ungirdled control and girdled, non-defoliated treatment, whereas different lowercase letters indicate significant differences among defoliation treatments within each disease category.

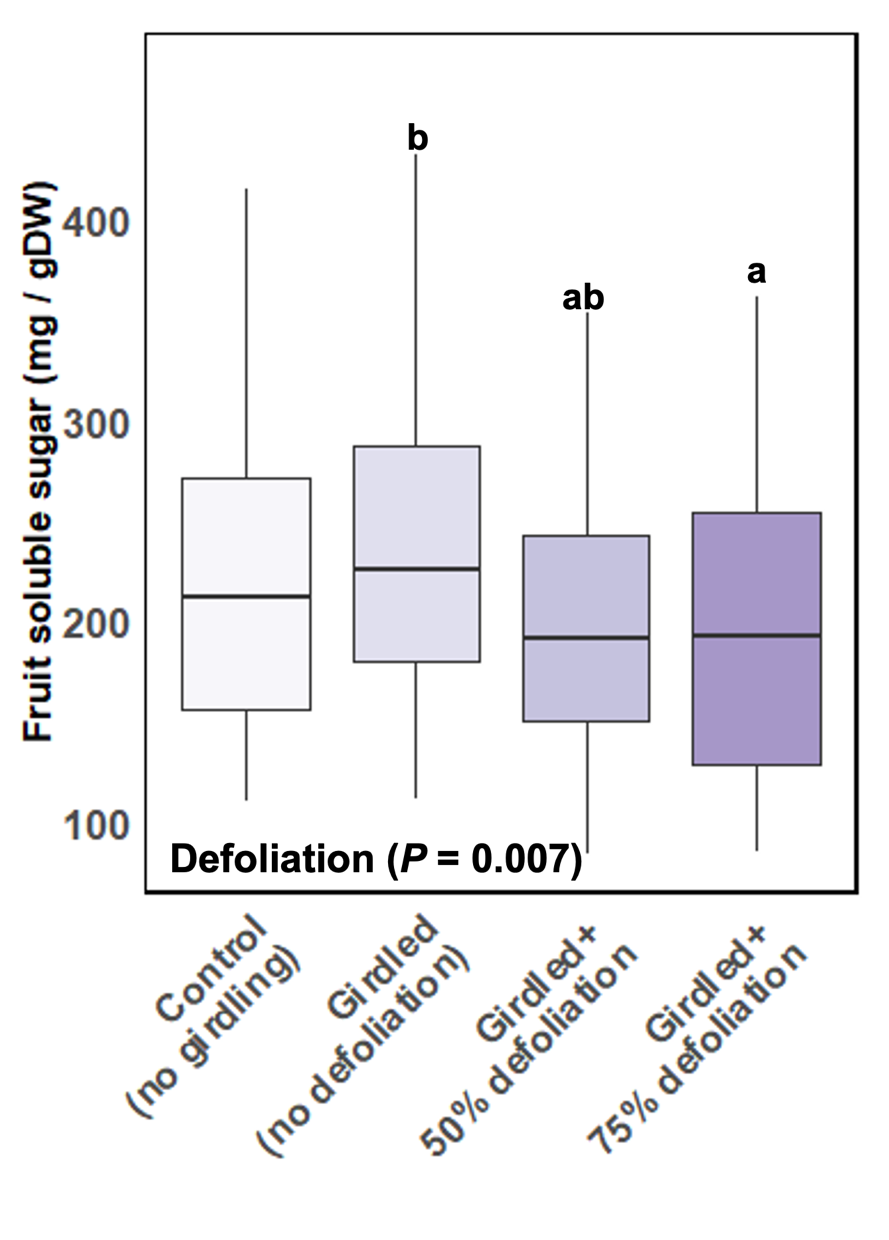

Figure S12**:** Effect of HLB disease on fruit soluble sugar level on *C. reticulata* ‘W. Murcott’ trees grown under field condition under protective screens at Stage II of fruit development. Source-sink manipulations were made on individual branches. Groups sharing different letters differ significantly (Tukey’s honestly significant difference test, *P* < 0.05). To distinguish girdling from defoliation effects, different capital letters indicate significant differences between the ungirdled control and girdled, non-defoliated treatment, whereas different lowercase letters indicate significant differences among defoliation treatments within each disease category.

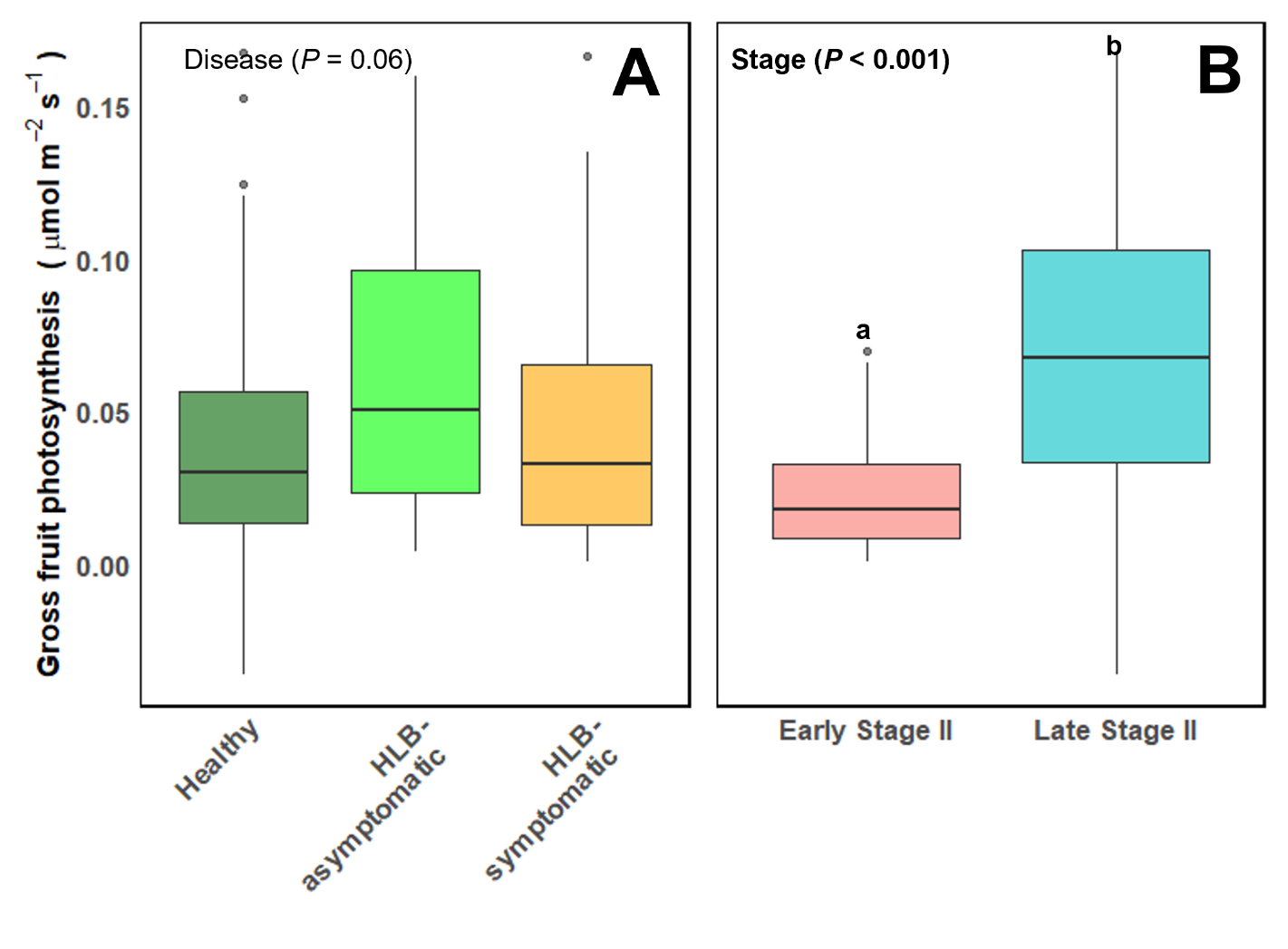

Figure S13**:** Fruit photosynthesis response of *C. reticulata* ‘W. Murcott’ grown under protective screens to HLB disease (A) and Fruit development (B) at Stage II of fruit development. Groups labeled with different letters are significantly different (Tukey’s Honestly Significant Difference; *P* < 0.05).

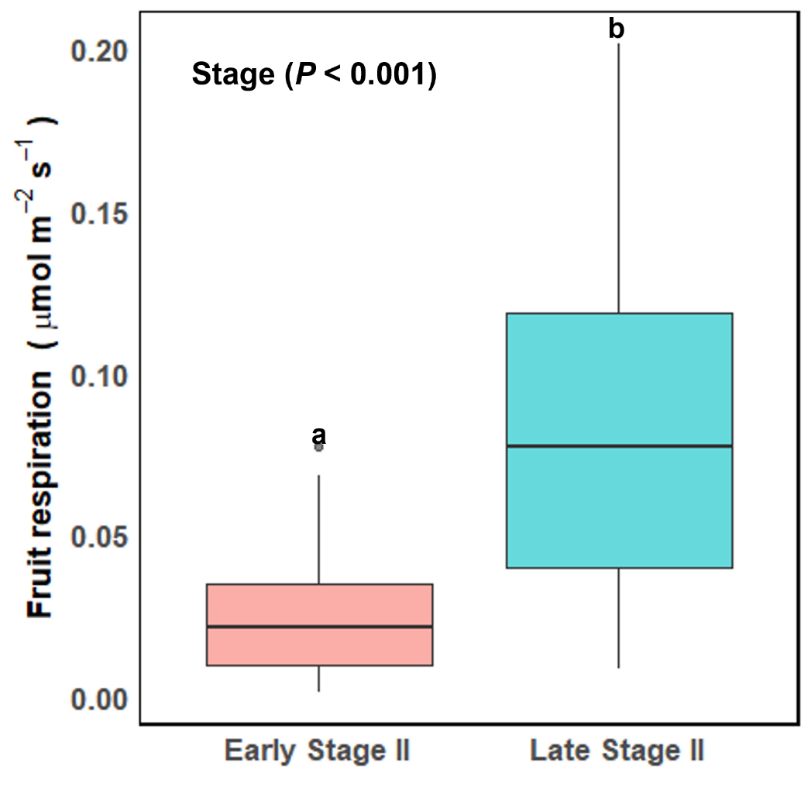

Figure S14**:** Fruit respiration at early and late Stage II of fruit development in *C. reticulata* ‘W. Murcott’ grown under filed condition under protective screens. Groups with different letters at significantly different (Tukey’s Honestly Significant Difference; *P* < 0.05).
