## Supplementary tables supporting results for "Phloem-Limited Disease Impacts Citrus Carbohydrate Dynamics Distinctly from Source-Sink Manipulation"

Table S1: Average Ct values of field grown healthy and HLB diseased *C. reticulata* ‘W. Murcott’ under protective screens used for the study.

| **Disease categories** | **Average Ct values** |
| --- | --- |
| Healthy | 35.4 ± 0.69 |
| HLB-asymptomatic | 30.4 ± 0.9 |
| HLB-symptomatic | 34.5 ± 1.16 |

Table S2: Analysis of variance for leaf gas exchange of field grown healthy and HLB diseased *C. reticulata* ‘W. Murcott’ under protective screens at varying source:sink ratio. HLB disease status of the trees were determined using qPCR assays and source-sink manipulations were made on individual branches. D- Disease status; T- Source-sink treatment; S- Stages of fruit development, *V_c_*_max_ - maximum carboxylation rate of rubisco, *J*_max_ - photosynthetic electron transport rate, *A*_leaf_ -leaf net photosynthesis, *Rd*_leaf_ – dark respiration, *Rlight*_leaf_ – daytime respiration, *L*_s_ -Stomatal limitation. The significant *P*-value was set at 0.05.

| **Variable** | **Effect** | **Num. DF** | **Den.DF** | **F value** | ***P* value** | **Variable** | **Num.**  **DF** | **Den. DF** | **F value** | ***P* value** |
| --- | --- | --- | --- | --- | --- | --- | --- | --- | --- | --- |
| ***V_c_*_max_** | **D** | 2 | 31.0 | 1.6 | 0.2 | ***J*_max_** | 2 | 27.2 | 1.6 | 0.2 |
|  | **T** | 3 | 88.3 | 1.5 | 0.2 |  | 3 | 65.2 | 0.9 | 0.4 |
|  | **S** | 1 | 100.0 | 2.2 | 0.1 |  | 1 | 76.9 | 0.9 | 0.3 |
|  | **D*T** | 6 | 87.5 | 0.3 | 0.9 |  | 6 | 65.2 | 0.4 | 0.8 |
|  | **D*S** | 2 | 98.2 | 2.2 | 0.1 |  | 2 | 75.5 | 0.6 | 0.5 |
|  | **T*S** | 3 | 88.4 | 0.4 | 0.7 |  | 3 | 65.4 | 1.2 | 0.3 |
|  | **D*T*S** | 6 | 87.2 | 1.1 | 0.3 |  | 6 | 64.9 | 2.0 | 0.07 |
| ***A*_net_** | **D** | 2 | 30.9 | 4.3 | 0.02 | ***R*_light_** | 2 | 23.6 | 1.2 | 0.3 |
|  | **T** | 3 | 88.3 | 2.1 | 0.09 |  | 3 | 87.6 | 2.2 | 0.09 |
|  | **S** | 1 | 99.9 | 0.2 | 0.6 |  | 1 | 97.3 | 13.9 | 0.0003 |
|  | **D*T** | 6 | 87.5 | 0.2 | 0.048 |  | 6 | 86.8 | 1.0 | 0.3 |
|  | **D*S** | 2 | 98.0 | 0.3 | 0.7 |  | 2 | 90.5 | 2.5 | 0.08 |
|  | **T*S** | 3 | 88.5 | 1.0 | 0.3 |  | 3 | 87.8 | 2.1 | 0.09 |
|  | **D*T*S** | 6 | 87.3 | 1.7 | 0.1 |  | 6 | 86.3 | 1.8 | 0.09 |
| ***R*_dark_** | **D** | 2 | 17.1 | 3.7 | 0.04 |  |  |  |  |  |
|  | **T** | 3 | 48.3 | 0.3 | 0.7 |  |  |  |  |  |
|  | **D*T** | 6 | 48.3 | 0.2 | 0.9 |  |  |  |  |  |
| ***L*_s_** | **D** | 2 | 27.2 | 0.54 | 0.5 |  |  |  |  |  |
|  | **T** | 3 | 64.5 | 7.89 | 0.0001 |  |  |  |  |  |
|  | **S** | 1 | 77.2 | 1.74 | 0.1 |  |  |  |  |  |
|  | **D*T** | 6 | 64.7 | 9.05 | <0.001 |  |  |  |  |  |
|  | **D*S** | 2 | 76 | 0.66 | 0.5 |  |  |  |  |  |
|  | **T*S** | 3 | 64.7 | 1.35 | 0.2 |  |  |  |  |  |
|  | **D*T*S** | 6 | 64.4 | 1.84 | 0.1 |  |  |  |  |  |

Table S3: Analysis of variance for leaf CO_2_ gas exchange of field grown on effect of disease on *C. reticulata* ‘W. Murcott’ under protective screens for Control and 75% defoliated treatment. HLB disease status of the trees were determined using qPCR assays and source-sink manipulations were made on individual branches. *A*_leaf_ -leaf net photosynthesis, *Rd*_leaf_ – dark respiration, *g*_s_ cost – stomatal cost, *L*_s_ -Stomatal limitation. The significant *P*-value was set at 0.05.

| **Variable** | **Treatment** | **Num. DF** | **Den.DF** | **F value** | ***P* value** |
| --- | --- | --- | --- | --- | --- |
| *g*_s_ cost | Control | 2 | 23.9 | 0.7 | 0.5 |
| *g*_s_ cost | 75% defoliation | 2 | 18.9 | 3.2 | 0.05 |
| *Rd*_leaf_ | Control | 2 | 104.3 | 31.2 | <0.0001 |
| *Rd*_leaf_ | 75% defoliation | 2 | 50.0 | 5.05 | 0.01 |
| *A*_leaf_ | Control | 2 | 30 | 4.9 | 0.01 |
| *A*_leaf_ | 75% defoliation | 2 | 19.1 | 6.5 | 0.006 |
| *L*_s_ | Control | 2 | 22 | 2.8 | 0.07 |
| *L*_s_ | 75% defoliation | 2 | 14.7 | 12.6 | 0.0006 |

Table S4: Analysis of variance for stem non-structural carbon of field grown healthy and HLB diseased *C. reticulata* ‘W. Murcott’ under field conditions under protective screens at Stage II of fruit development and varying source:sink ratio. HLB disease status of the trees were determined using qPCR assays and source-sink manipulations were made on individual branches. D- HLB Disease status; T- Source-sink treatment; S- Stage of fruit development. The significant *P*-value was set at 0.05.

| **Variable** | **Effect** | **Num. DF** | **Den.DF** | **F value** | ***P* value** | **Variable** | **Num.**  **DF** | **Den. DF** | **F value** | ***P* value** |
| --- | --- | --- | --- | --- | --- | --- | --- | --- | --- | --- |
| **Stem sucrose** | **D** | 2 | 33.4 | 2.2 | 0.1 | **Stem starch** | 2 | 27.8 | 1.0 | 0.3 |
|  | **T** | 3 | 105.6 | 6.2 | <0.001 |  | 3 | 106.3 | 15.2 | <0.001 |
|  | **S** | 1 | 126.6 | 3.1 | 0.07 |  | 1 | 124.5 | 60.4 | <0.001 |
|  | **D*T** | 6 | 105.6 | 2.4 | 0.02 |  | 6 | 106.3 | 3.4 | 0.003 |
|  | **D*S** | 2 | 126.8 | 3.9 | 0.02 |  | 2 | 121.2 | 0.01 | 0.9 |
|  | **T*S** | 3 | 105.6 | 1.0 | 0.3 |  | 3 | 107 | 0.6 | 0.5 |
|  | **D*T*S** | 6 | 105.6 | 1.1 | 0.3 |  | 6 | 106 | 1.3 | 0.2 |
| **Stem soluble sugar** | **D** | 2 | 24.03 | 3.3 | 0.05 |  |  |  |  |  |
|  | **T** | 3 | 106. | 1.82 | 0.1 |  |  |  |  |  |
|  | **S** | 1 | 126.4 | 276.7 | <0.001 |  |  |  |  |  |
|  | **D*T** | 6 | 106. | 1.36 | 0.2 |  |  |  |  |  |
|  | **D*S** | 2 | 121.5 | 1.94 | 0.1 |  |  |  |  |  |
|  | **T*S** | 3 | 106.1 | 0.56 | 0.6 |  |  |  |  |  |
|  | **D*T*S** | 6 | 106.1 | 0.9 | 0.4 |  |  |  |  |  |

Table S5: Analysis of variance for fruit growth, fruit gas exchange and non-structural carbon of fruits of field grown healthy and HLB diseased *C. reticulata* ‘W. Murcott’ under protective screens at varying source:sink ratio and Stage II of fruit development. HLB disease status of the trees were determined using qPCR assays and source-sink manipulations were made on individual branches. D- HLB Disease status; T- Source-sink treatment; S- Stage of fruit development. The significant *P*-value was set at 0.05.

| **Variable** | **Effect** | **Num. DF** | **Den.DF** | **F value** | ***P* value** | **Variable** | **Num.**  **DF** | **Den. DF** | **F value** | ***P***  **value** |
| --- | --- | --- | --- | --- | --- | --- | --- | --- | --- | --- |
| **Fruit DW** | **D** | 2 | 29.4 | 2.3 | 0.1 | **Fruit diam** | 2 | 31.0 | 0.1 | 0.8 |
|  | **T** | 3 | 100.2 | 2.8 | 0.04 |  | 3 | 106.7 | 4.5 | 0.004 |
|  | **S** | 1 | 123.36 | 140.5 | <0.001 |  | 1 | 131 | 20.1 | <0.001 |
|  | **D*T** | 6 | 100.2 | 0.7 | 0.6 |  | 6 | 106.7 | 0.3 | 0.9 |
|  | **D*S** | 2 | 100.2 | 0.5 | 0.6 |  | 2 | 131.1 | 0.08 | 0.9 |
|  | **T*S** | 3 | 123.6 | 4.3 | 0.01 |  | 3 | 106.7 | 0.06 | 0.9 |
|  | **D*T*S** | 6 | 100.2 | 0.3 | 0.8 |  | 6 | 106.7 | 1.3 | 0.2 |
| ***A*_net_** | **D** | 2 | 23.6 | 3.1 | 0.06 | ***R*_dark_** | 2 | 124 | 1.5 | 0.2 |
|  | **T** | 3 | 99.6 | 4.4 | 0.005 |  | 3 | 124 | 2.5 | 0.05 |
|  | **S** | 1 | 126.8 | 77.8 | <0.001 |  | 1 | 124 | 32.3 | <0.001 |
|  | **D*T** | 6 | 99.6 | 0.4 | 0.8 |  | 6 | 124 | 0.7 | 0.6 |
|  | **D*S** | 2 | 126.9 | 0.7 | 0.4 |  | 2 | 124 | 1.0 | 0.3 |
|  | **T*S** | 3 | 99.6 | 0.1 | 0.9 |  | 3 | 124 | 0.8 | 0.4 |
|  | **D*T*S** | 6 | 99.6 | 1.4 | 0.1 |  | 6 | 124 | 0.6 | 0.6 |
| **Fruit sucrose** | **D** | 2 | 25.2 | 2.5 | 0.1 | **Fruit starch** | 2 | 21.3 | 30.1 | <0.001 |
|  | **T** | 3 | 102.7 | 2.7 | 0.04 |  | 3 | 97.5 | 3.1 | 0.02 |
|  | **S** | 1 | 124.8 | 24.4 | <0.001 |  | 1 | 117 | 52.1 | <0.001 |
|  | **D*T** | 6 | 102.7 | 0.6 | 0.6 |  | 6 | 97 | 1.2 | 0.2 |
|  | **D*S** | 2 | 122.9 | 1.3 | 0.2 |  | 2 | 110 | 2.3 | 0.1 |
|  | **T*S** | 3 | 102.8 | 1.1 | 0.3 |  | 3 | 97 | 0.4 | 0.7 |
|  | **D*T*S** | 6 | 102.8 | 1.1 | 0.9 |  | 6 | 97 | 1.1 | 0.3 |
| **Fruit soluble sugar** | **D** | **2** | 26.3 | 0.2 | 0.8 |  |  |  |  |  |
|  | **T** | 3 | 99.5 | 4.23 | 0.007 |  |  |  |  |  |
|  | **S** | 1 | 125.5 | 127.1 | <0.001 |  |  |  |  |  |
|  | **D*T** | 6 | 99.5 | 0.45 | 0.8 |  |  |  |  |  |
|  | **D*S** | 2 | 125.8 | 2.32 | 0.1 |  |  |  |  |  |
|  | **T*S** | 3 | 99.6 | 1.89 | 0.1 |  |  |  |  |  |
|  | **D*T*S** | 6 | 99.7 | 0.85 | 0.5 |  |  |  |  |  |
